# The apoplastic space of two wheat genotypes provide highly different environment for pathogen colonization: Insights from proteome and microbiome profiling

**DOI:** 10.1101/2023.06.05.543792

**Authors:** Carolina Sardinha Francisco, Mohammad Abukhalaf, Clara Igelmann, Johanna Gustke, Michael Habig, Liam Cassidy, Andreas Tholey, Eva Holtgrewe Stukenbrock

## Abstract

The intercellular space comprising the plant apoplast harbors a diverse range of microorganisms. The apoplastic interface represents the main compartment for interactions between proteins produced and secreted by the plant and the microbial endophytes. The outcomes of these interactions can play a role in plant cell wall metabolism, stress tolerance, and plant-pathogen resistance. So far the underlying factors that determine microbiota composition in the apoplast are not fully understood. However, it is considered that cell wall composition, nutrient availability, and the plant immune system are main determinants of microbiota composition. The plant immune system is considered to play a crucial role in modulating microbiota composition through the recognition of specific microbe-associated molecular patterns and the activation of defense responses. Hereby the plant may restrict non-beneficial microbial members and facilitate the propagation of beneficial ones. In this study, we investigated changes in the apoplastic environment during pathogen invasion using wheat as a model system. Infection of wheat with Zymoseptoria tritici, a fungal pathogen, resulted in notable alterations in the apoplast composition, reduced microbial diversity, and the accumulation of antimicrobial defense metabolites. Intriguingly, certain core microbial members persisted even in the presence of pathogen-induced immune responses, indicating their ability to evade or tolerate host immune defenses. To further explore these dynamics, we developed a protocol for extracting apoplastic fluids from wheat leaves and conducted proteome analyses to characterize the dynamic environment of the wheat leaves. Our findings uncovered a highly variable apoplastic environment that selects for microbes with specific adaptations. Notably, a core microbial community enriched in the resistant wheat cultivar exhibited antagonistic activity against Z. tritici, suggesting a potential role in conferring pathogen defense. This study advances our understanding of the dynamic interactions and adaptations of the wheat apoplastic microbiota during pathogen invasion, emphasizing the pivotal role of microbial interactions in pathogen defenses.

## Introduction

Plants are hosts to complex communities of microorganisms that function together in symbiotic relationships that define the “plant metaorganism” (1, 2). Detailed studies of microbial communities in below and above ground tissues have demonstrated that different plant compartments host distinct microbial communities (reviewed in (3)). A large diversity of microorganisms colonizes the surface of plant tissues, while some microorganisms have adapted to colonize the interior of plant tissues. Hereby, also the apoplastic compartment, which is the intercellular space within plant tissues, hosts a diverse range of microorganisms, including both bacteria and fungi. The composition of the apoplastic microbiota can vary depending on the tissue type and the developmental stage of the plant, but it is considered to play a role in plant cell wall metabolism and stress tolerance, including pathogen resistance (4, 5).

As distinct plant compartments represent very different environments for microbial propagation, they may select for microbes with particular adaptations. In the apoplastic compartment, factors which may select for particular microbial partners include cell wall composition, nutrient availability and the host immune system. For example, experimental studies using mutants of *Arabidopsis* affected in cuticle synthesis showed a pronounced influence on microbiota composition (6). Moreover, the apoplast is generally a nutrient-poor environment, and therefore favors microorganisms that can utilize complex plant compounds and effectively scavenge nutrients (7). Importantly, the apoplast is the main compartment for interactions between proteins produced and secreted by the plant and the microbial endophytes. How these factors jointly shape the plant microbiota during the lifetime of a plant remains poorly understood. Yet, understanding how the apoplastic environment shapes microbial diversity is crucial to our understanding of plants as metaorganisms as well as to the development of synthetic microbial communities intended to strengthen plant health in production systems.

Genetic and experimental studies of plant-microbiota interactions have demonstrated a fundamental role of the plant immune system in shaping microbiota composition. For example, mutants of *A. thaliana* impaired in different immune-related signaling pathways showed altered composition of their microbiota (e.g., (8, 9). Likewise, activation of systemic immune responses in wheat, *Triticum aestivum*, by an avirulent pathogen, conferred a significant shift in microbiota composition correlating with the production of defense related metabolites (10). The evident function of the plant immune system in shaping microbiota composition leads to the general assumption that plants, via immune signaling, can regulate microbiota composition by selecting for beneficial microbial partners and eliminating non-beneficial, pathogenic microbes. Conceptually, these different types of microbial regulations have been termed “recognition and tolerance" and "recognition and defense" responses (3, 11). At the core of these interactions are pattern recognition receptors (PRR proteins) which are embedded in plant membranes and act to recognize and distinguish particular microbe associated molecular patterns (MAMPs) produced by microorganisms (12, 13). The activation of defense signaling involves the production of reactive oxygen species (ROS) and specific antimicrobial metabolites, modification of cell wall compositions, and eventually local cell death. Such "recognition and defense responses” may lead to the elimination of certain microorganisms, while simultaneously selecting for other microorganisms that can tolerate a higher abundance of plant defense compounds (9). This scenario invokes that core members of the plant metaorganism have adapted to tolerate defense-related plant compounds, which are recurrently induced by invading non-beneficial and pathogenic microorganisms.

In this study, we set out to investigate the plant apoplastic space as an environment for microbial colonization. We specifically addressed how induced immune responses alter the abiotic and biotic composition of the apoplast. Furthermore, we investigated signatures of microbial adaptation to the apoplastic environment by growth assays of plant-associated microbes in the presence of plant immune compounds. We used wheat as a model system to dissect changes in apoplastic fluid compositions. Wheat is one of the most important crops grown world-wide. In recent years, research has focused on the wheat microbiome considering the putative potential of microbiota- manipulation in plant growth promotion and stress tolerance (e.g., (14–16). In this regard, insights into the dynamics of microbiota composition are scarce, hindering our ability to predict the stability of beneficial microbes in the field.

In our study, we used wheat as a plant model system to uncover variation in the apoplast environment. Wheat can be infected by the ascomycete fungus *Zymoseptoria tritici*, a hemibiotrophic pathogen that invades leaf tissues via stomatal openings. Several quantitative and qualitative resistance loci have been mapped in wheat using quantitative trait locus analyses and genome wide association studies (17–21). Among these resistance loci the best characterized is *Stb6*, which confers resistance towards *Z. tritici* isolates by the secretion of the AvrStb6 protein (17, 22, 23). In resistant wheat varieties harboring *Stb6*, the recognition of AvrStb6 in *Z. tritici* isolates occurs via guard-cell located in the stomata, preventing fungal invasion and colonization (24). Hereby, recognition of *Z. tritici* induces a strong up-regulation of local and systemic defense responses involving the production of ROS and other defense related compounds, as well as lignin accumulation (10). In susceptible wheat however, the fungus efficiently suppresses defense signaling to colonize the mesophyll tissue. In a previous study, we compared the impact of microbiota composition in wheat leaves after inoculation with spores of *Z. tritici* (10). We showed that the fungal pathogen greatly influences microbiota composition in both resistant and susceptible wheat. We speculate that the microbiota composition may change in response to both direct interactions with the fungus as well as indirect effects conferred by the altered apoplast environment.

An interesting observation is the fact that microbial diversity overall is reduced in a resistant wheat cultivar upon infection with *Z. tritici* (10). The reduced microbial diversity correlates with a strong accumulation of antimicrobial defense metabolites which are induced by the invading pathogen. One component of the microbial community however persists during the pathogen-induced immune response suggesting that these microbes are capable of avoiding or tolerating host immune responses.

In the present study, we set out to further explore the dynamic of abiotic and biotic conditions in a wheat metaorganism. We notably considered changes in the apoplastic space during pathogen invasion and explored the diversity and adaptations of core metaorganisms members. To this end, we developed a protocol for extraction of apoplastic fluids from wheat leaves. We used the apoplastic fluid samples for proteome analyses and for isolation of the microbial communities inhabiting the particular intercellular space of the wheat mesophyll.

Our study reveals a highly variable environment, selecting for microbes with particular adaptations. We observed a portion of microbes that are specifically enriched in the resistant cultivar Chinese Spring (which harbors *Stb6*), and we hypothesize that they compose a core microbial community. We further observed that several of these core microbes show antagonism against *Z. tritici* and may therefore also play a role in pathogen defense.

## Material and methods

### Fungal and plant material

The Dutch *Z. tritici* IPO323 (internally named Zt244) strain producing the effector protein AvrStb6 was used in this study (Wageningen University, Netherlands) (17, 22, 25). Additionally, the Zt05 strain isolated in Denmark (26), and Zt10 strain isolated in Iran (27), were used in the *in-vitro* confrontation assays. The fungal cells, stored in glycerol at -80°C, were recovered in 50 mL of YMS medium (yeast malt sucrose broth, 4 g/L yeast extract, 4 g/L malt extract, 4 g/L sucrose, pH 6.8) incubated at 18 °C for 4 days. The susceptible wheat (*Triticum aestivum* L) cultivar Obelisk (without the resistance gene *Stb6*) was obtained from Wiersum Plant Breeding BV (the Netherlands), and the resistant landrace Chinese Spring (with the resistance gene *Stb6*) was kindly provided by Bruce McDonald from ETH Zürich.

### Plant growth conditions

Seeds from both Obelisk and Chinese Spring wheat genotypes were surface washed for 5 min with 1x phosphate-buffered saline (PBS; 8 g/L NaCl, 0.2 g/L KCl, 1.44 g/L Na2HPO4, 0.24 g/L KH2PO4, pH 7.4) supplemented with 0.02% (v/v) of Tween 20, followed by three times (1 min each) with sterile deionized water. Twelve seeds of each wheat genotype were sown in square pots (9 x 9 x 9.5 cm) containing peat soil (HAWITA GmbH, Vechta, Germany). Subsequently, each pot was inoculated with 100 mL of agricultural soil slurry by diluting 10 g of soil collected from the experimental farm Hohenschulen (54°18’53.7” N – 9°58’44.1” E) of the Christian-Albrechts University of Kiel in Germany in 1L of sterile deionized water. Plants were grown during the entire experiment in the greenhouse under constant conditions of 16/8 h light/dark cycle at 22°C (day) and 20°C (night) and 80% relative humidity.

### Fungal infection

Plant inoculations were performed as previously described (28). In brief, the IPO323 strain was grown in 50 mL of YMS medium in a rotary shaker for five days at 18°C. After the incubation, blastospore suspensions were filtered through sterile cheesecloth and pelleted at 3750 rpm for 15 min at 4°C. The supernatant was discarded and the cells were resuspended in sterile deionized water. The concentration of the blastospore suspensions were determined using KOVA Glasstic counting chambers (KOVA International Inc., USA) and adjusted to a final concentration of 10^7^ blastospores/mL in 30 mL of sterile water supplemented with 0.1% (v/v) Tween 20. We sprayed 14-day old plants until run-off and placed for three days in sealed plastic bags to keep humidity at 100%. The plastic bags were then removed, and the plants were kept in the greenhouse until the end of the experiments. Mock-treated plants were sprayed with sterile water. To ensure the success of fungal infection, second leaves of additional infected plants were harvested at 14 days-post inoculation (dpi).

### Wheat apoplastic fluids extraction

Apoplastic fluids (AFs) was obtained according to the method of R. T. Nakano et al. (29) with slight modifications (Fig. S1A). Ten 1^st^ and 2^nd^ fully expanded wheat leaves were harvested at 8 dpi from plants growing in separated pots. Three biological replicates containing 20 leaves (the 10 first and 10 second wheat leaves) were pooled for the AF extraction. Leaves were surface washed two times (5 min each time) with 1x PBS, and 1x PBS supplemented with 0.02% (v/v) of Tween 20, followed by three times (1 min each) with sterile deionized water. The washed leaves were submerged in 400 mL of infiltration buffer (5 mM sodium acetate, 0.2 M calcium chloride, pH 4.3) containing two dissolved cOmplete^TM^ EDTA-free protease inhibitor cocktail tablets (Roche Applied Science, Penzberg, Germany) and vacuum-infiltrated at 900 mbar for 10 min. Subsequently, the vacuum was released gently for approximately 30 min. The vacuum step was carried out 2-3 times to reach a maximum saturation of the infiltration buffer in the apoplastic space. The excessive buffer was dried quickly between two sheets of soft paper towel, and the leaves were placed into a 20 mL blunt-end needleless syringe (without the plunger). The syringe was introduced into a 50 mL falcon tube and then centrifuged at 4000 rpm for 4 minutes at 4 °C. The centrifugation step was repeated up to three times to extract a maximum amount of the apoplastic liquid. A summary of the methodology used for the apoplastic fluid isolation is given in Fig. S1A. Two 500 µL aliquots of AFs were lyophilised overnight, followed by resuspension in 200 µL of 100 mM ammonium acetate, and then stored at -20° until required for proteomic or Western blot analyses. Additionally, 1 mL of AFs were kept on ice until plated on different media for microbial isolation.

### Western blot analysis

Western blotting was performed with three biological replicates of each treatment to identify any contamination of the apoplastic fluid samples by intracellular proteins. The cytoplasmic protein Ribulose-1,5-bisphosphate carboxylase/oxygenase (so-called RubisCO, RbcL) was used as a marker for intra-cellular contamination of the apoplastic fluid samples. Since different eukaryotes colonize the apoplast, including *Z. tritici* for the wheat infected plants, we furthermore used the Histone H3 protein as a general eukaryote positive marker. One microgram of protein from the apoplastic fluids and a quantitative rubisco positive standard control (Agrisera, Vännäs, Sweden) were electrophoretic separated on Mini-PROTEAN TGX Precast Gels (8-16% polyacrylamide, Bio-Rad Laboratories, California, USA) at 70V for 15 min, and then increased to 100V for 60 min, followed by blotting to PVDF membranes (Bio-Rad Laboratories, California, USA). Membranes were overnight blocked with 1x TBS-T (Tris-buffered saline with Tween-20) containing 4% of non-fat dry milk at 4°C with agitation. Blots were then incubated in the primary antibodies against Histone H3 (H3, Agrisera, Vännäs, Sweden) (1:5’000) or RuBisCo large subunit (RbcL, Agrisera, Vännäs, Sweden) (1:10’000) for 1h at room temperature with agitation. After washing, the blots were incubated in 1x TBS-T containing 2% of non-fat dry milk in which the secondary antibodies conjugated to horseradish peroxidase (anti-rabbit, 1:20’000 from Santa Cruz Biotechnology, Dallas, USA; or anti-chicken, 1:25’000 from Agrisera, Vännäs, Sweden) for 1h at room temperature. The blots were detected using ECL-Select Western Blotting detection reagent (Cytiva, München, Germany) according to the manufacture’s instruction, and images were obtained using the CCD imager Documentation System (Bio-Rad Laboratories, California, USA).

### Proteome analysis by liquid chromatography-mass spectrometry (LC-MS)

Protein concentrations in the apoplast samples were determined using the micro BCA assay (Thermo Fischer Scientific, Germany). 50 µg total protein of each sample were reduced with 10 mM dithiothreitol (DTT) at final concentration for 1h at 56°C. Samples were then alkylated with 50 mM Chloroacetamide (CAA) for 30 min at 25°C in the dark, followed by overnight digestion with trypsin (enzyme to protein ratio of 1:100) (Promega, Germany) at 37 °C. Digestion was stopped by acidification using 5% formic acid (FA) and samples were subsequently desalted on a C18 solid phase extraction cartridge (SepPak C18, 1cc, 50 mg) (Waters, Eschborn, Germany) following manufacturer’s instructions. Peptides were eluted from the cartridges with 60% acetonitrile (ACN) supplemented with 0.1% trifluoroacetic acid (TFA), and dried via vacuum centrifugation (Concentrator Plus, Eppendorf).

Dried peptides were resuspended in 3% ACN and 0.1% TFA and then separated employing a Dionex U3000 nanoHPLC system (Dreieich, Germany) equipped with an Acclaim Pepmap100 C18 column (2 µm, 75 µm x 500 mm). The eluents used were eluent A: 0.05% FA, and eluent B: 80% ACN/ 0.04% FA. A programmed 123 min run at 300 nl/min was performed as follows: 2 min at 4% eluent B, 90 min linear gradient 4% to 50% eluent B, 5 min linear gradient 50% to 90% eluent B, 10 min at 90% eluent B, followed by equilibration for 16 min at 4% eluent B. Eluted peptides were analyzed online on a QExactive^TM^ Plus mass spectrometer (Thermo, Bremen, Germany). A top15 data acquisition method was applied with a full MS scan (350-1400 m/z, resolution 70,000, AGC 3e6, Max IT 50 ms), followed by HCD fragmentation and MS/MS scans of the 15 most intense ions (NCE 27.5, resolution 17,500, AGC 1e5, Max IT 100 ms). A 20 s dynamic exclusion and a lock mass of 445.12003 were applied. Each sample was injected twice.

### Proteomic data analysis

MS raw data were analyzed using the Uniprot reference proteomes of *Triticum aestivum* (130,673 proteins), *Z. tritici* strain IPO323 (10,972), *Acidovorax sp*. (5,416), *Curtobacterium flaccumfaciens* (3,574), *Pseudomonas anguilliseptica* (4,385), *Pseudomonas rhodesiae* (5,708), *Flavobacterium sp* (3,029) *Plantibacter flavus* (3,813), and known contaminants (cRAP) were determined using the Sequest HT search engine linked to Proteome Discoverer^TM^ 2.5 (Thermo Fischer Scientific, Germany). Enzyme specific was set to Trypsin, and two missed cleavages were allowed. Precursor and fragment ion tolerances were set to 10 ppm and 0.02 Da, respectively. Carbamidomethylation of cysteine was set as a fixed modification and oxidation of methionine as a variable modification. The INFERYS rescoring node was incorporated into the processing workflow. Strict parsimony criteria were applied with 1% false discovery rate (FDR) cutoff used for both peptides and proteins. A label-free quantification method, based on the intensities of the precursor ions, was performed using Minora detection with match between runs. Only proteins with high combined FDR confidence were considered. Moreover, *T. aestivum* proteins were filtered to contain at least two identified peptides; proteins of other organisms were considered underrepresented in the samples.

Protein groups and their respective raw intensities were exported and further analyzed using the software Perseus v.1.6.15.0 (30). For quantitative analysis, intensities of two technical injections per sample were averaged and then normalized by median based normalization. Normalized intensities were log_2_ transformed and grouped per condition (*e.g*., Obelisk Mock or Chinese Spring Mock). Proteins were filtered to contain at least three values in one group, and imputation of missing values from normal distribution was performed separately for each column (Perseus default values, Width 0.3 and down shift 1.8). Principal component analysis (PCA) plots were generated in Perseus and the volcano graphs were plotted with bioinfokit (31) using a custom made Python script.

### Gene enrichment analysis and predictions

Gene ontology (GO) enrichment analysis was performed with the webserver DAVID 2021 (32, 33). All of the identified wheat proteins were searched against the default background and functional annotation charts with GOTERM_BP_DIRECT, GOTERM_CC_DIRECT and GOTERM_MF_DIRECT categories (EASE 0.1). Up to five GO terms with lowest Benjamini Hochberg corrected p-values (<0.05) were plotted. GO enrichment of the differentially expressed proteins in the volcano plots were performed in a similar way; however, identified proteins were used as a background and up to three GO terms with lowest Benjamini Hochberg p-values (<0.05) were plotted for each category.

For prediction of apoplastic proteins, a FASTA file of the identified proteins was analyzed using AploplastP 1.0 (34) and SignalP 6.0 (Organism: Eukarya, Model mode: Fast) (35). Proteins predicted to be apoplastic by the program ApoplastP or having a signal peptide predicted by the SignalP software were considered as predicted apoplastic proteins. Venn diagrams were plotted using Venny 2.1 program (36).

### Establishment of a wheat-enriched endophytic community-based culture collection

Using the apoplastic fluid samples, we further isolated a collection of bacteria and fungi. The culture collection represents the microbial community which is enriched in the inter-cellular space of wheat leaves. The culture collection was obtained using freshly extracted AFs from mock and IPO323 infected Obelisk or Chinese Spring wheat plants. To optimize the isolation of microorganisms we used a set of different media. 100 µL of both undiluted or 1:10 dilution of AF samples were plated on three different media, including as Trypticase Soy agar (TSA; 7.5 g/L BBL Trypticase Soy Broth (TSB); Becton Dickinson and Company, NJ, USA; 15 g/L agar) used as a non-selective growth medium for bacteria; Reasoner’s 2A agar (R2A; 18.1 g/L R2A-Agar; Carl Roth, Karlsruhe, Germany) a medium that favors slow-growing bacteria; and *Pseudomonas syringae* medium (PSM; 15 g/L sucrose, 3.75 g/L peptone, 0.38 g/L K2HPO4, 0.1 g/L MgSO4, 15 g/L agar) recommended to isolate *Pseudomonas* species. Three technical replicate plates were prepared per treatment and media combinations. Plates were incubated for four days at 22°C or until the appearance of visible fastidious colonies. For microbial singularization, individuals exhibiting diversity colony morphotypes (*e.g.,* distinct size, shape, texture, or pigmentation) were transferred onto a new agar plate and incubated for additionally four days at 22°C. This procedure was repeated twice in at similar way. The individualized colonies were then transferred to canonical tubes containing 25 mL of TSB medium and incubated on a rotary shaker at 145 rpm for seven days at 22°C. Finally, cryostocks were prepared by mixing 1 mL of the liquid cultures with 1 mL of 50% (v/v) YMS glycerol medium.

### DNA extraction, species identification and phylogenetic analysis

Cells of individualized colonies were scratched from the agar and dissolved in 250 μL of sterile deionized water. 50 μL of the cell suspensions were lysed by adding 100 μL of TE lysis buffer (1 mM EDTA and 10 mM TrisHCl, pH 8 + 0.1 % Triton X 100) and incubated at 99 °C for 10 min. Subsequently, the samples were centrifuged at 13’000 *x g* for 10 min. The supernatant was transferred to a new tube and used as DNA template for PCR amplification. In order to identify the bacterial or fungal species, PCRs using the partial 16S rRNA and the ITS (internal transcribed spacer) regions were conducted, respectively (37, 38). The 16S rRNA V3-V7 gene region was PCR-amplified (94°C/2 min; 94°C/30 sec; 57°C/30 sec; and 72°C/1 min for 25 cycles, and final extension 72°C/10 min) using the forward primer 341F (5’ - CCT ACG GGA GGC AGC) and the reverse primer 1192R (5’ - ACG TCA TCC CCA CCT). Additionally, the ITS region was PCR-amplified (95°C/2 min; 95°C/30 sec; 55°C/30 sec; and 72°C/20 sec for 25 cycles, and final extension 72°C/10 min) using the forward primer ITS1F (5’- CTT GGT CAT TTA GAG GAA GT) and the reverse primer ITS1R (5’ - GCT GCG TTC TTC ATC GAT GC). The PCR products were purified using the Wizard^TM^ SV Gel and PCR Clean-Up System (Promega, Madison, WI, USA) following the manufactures instructions. The obtained purified PCR products were submitted for Sanger sequencing (Institute of Clinical Molecular Biology, Kiel, Germany) using the abovementioned forward primer sequences. To identify the microbial species, the nucleotide sequences were analyzed using the SILVA database with the incorporated SILVA ACT service (39) for homology searches, sequence alignment and taxa identification. Furthermore, individual sequences were blasted against the NCBI (National Center for Biotechnology Information) database. The nucleotide sequences were aligned using MEGA software (40). The best-fit model of nucleotide evolution was the GTR+G also determined by MEGA. The phylogeny reconstruction was computed by maximum likelihood (ML) using 500 bootstraps using RAxML program (41).

### Jaccard similarity index and phylogenetic analysis

The β-diversity (variation between treatment groups) was measured using the non-phylogenetic-based approach Jaccard index (42). Jaccard is a qualitative index that utilizes the presence-absence data of the species, as follows: the *S_i_*, *S_j_*, and *S_ij_* denote the number of species (OTUs) present in Samples *i*, *j*, and both *i* and j samples, respectively, the Jaccard distance between Sample *i* and *j* is represented as: 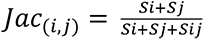. Distance matrices were constructed based on the percentage of dissimilarity of community-based culture collection.

A dataset containing the nucleotide sequences of all identified species was used for phylogenetic analysis. The two fungal species *Rhizopus arrhizus* and *Mucor kunryamgriensis* from the Mucoroycota were used as outgroups. The best-fit model of nucleotide evolution was the GTR+G model determined by MEGA v.11 (40). The nucleotide alignment was performed in MEGA v.11 using the Clustal W multiple sequence alignment (43), followed by a Maximum Likelihood method for phylogenetic reconstruction using 500 bootstraps performed in the RAxML v.8 (41, 44). The graphic display of the phylogenetic tree was adapted with the online tool iTOL v.5 (45).

### Confrontation assays

The potential antagonistic activities among the wheat-enriched endophytic microbes and *Z. tritici* were accessed by confrontation of co-cultures on Petri dishes. Considering the extensive genetic variation in *Z. tritici*, we included additionally two *Z. tritici* isolates in the assay allowing us to identify general and isolate-specific interactions of *Z. tritici* with wheat microbiome members. In addition to the Dutch isolate IPO323, we included two additional field isolates of *Z. tritici;* Zt05 from Denmark, and Zt10 from Iran both isolated from wheat (46).

To evaluate the antagonistic activity of the wheat-associated microbes against *Z. tritici*, the blastosporulation of the three tested *Z. tritici* strains was recovered on 50 mL of YMS liquid medium incubated at 18°C for six days. The 20 mL of soft agar medium was prepared using 15 mL of sterile deionized water, 5 mL of boiling Fries 3 agar medium supplemented with trace elements (as previously described), and fungal inoculum which gives a final concentration of 1.88x10^5^ blastospores/mL. The wheat- associated microbes were then recovered in Fries 3 liquid medium for three or four days before the assays on a rotary shaker (200 rpm) at 23°C. Microbial cell densities were measured and adjusted to OD^600^ of 0.2 at 600 nm. Finally, drops of 3.5 µL of each tested wheat endophyte were spotted onto three independent Fries 3 plates amended with *Z. tritici* spores. Plates containing solely Fries 3 medium were used as control. Nine different isolates placed 3.5 cm apart were tested per Petri dish. Each isolate was tested three times on independent plates. 1.5 µL of Hygromycin B (at a final concentration of 19.61 mg/L) was used as positive control for fungal growth inhibition. Plates were incubated in the dark at 23 °C and photographed after three or four days of incubation. Digital images were processed using the ImageJ software (47) for the quantification of the inhibition zones. A Kruskal-Wallis test was performed to determine the differences between the size of inhibition for the three tested *Z. tritici* strains. The Dunn’s test was then used to define the significance difference between each independent group. The association between the percentage of isolates that shown growth promotion or inhibition by each tested *Z. tritici* strain and the treatment from where the wheat-associated microbes were isolated was examined using Fisher’s exact test.

### Response to plant immune-related compounds

To investigate adaptations of the endophyte to the apoplastic environment, we conducted an *in vitro* assay with different plant-stress related molecules. We included both bacteria and basidiomycete yeasts in the assay. Hydrogen peroxide (H_2_O_2_) and three different secondary metabolites, including cinnamyl alcohol (CA), quinic acid (QA), and nonanoic acid (NA) were used to determine the tolerance of the wheat-associated microbes to plant immune responses. H_2_O_2_ accumulates in the apoplast and is widely used to mimic ROS burst upon Microbe-Associated Molecular Patterns (MAMPs) perception (48): CA, QA, and NA were previously demonstrated to be highly abundant in the Chinese Spring, but not in Obelisk upon *Z. tritici* infection (49). We consider these metabolites to be defense-related metabolites which are induced specifically in *Z. tritici* resistant wheat. To test for microbial tolerance, the microbes were recovered in Fries 3 liquid medium (5 g/L C_4_H_12_N_2_O_6_, 1 g/L NH_4_NO_3_, 0.5 g/L MgSO_4_*7H_2_O, 1.3 g/L KH_2_PO_4_, 2.6 g/L K_2_HPO_4_, 1 g/L yeast extract, 30 g/L sucrose) for 24h on a rotary shaker (200 rpm) at 23°C. Microbial cell densities were measured and adjusted to OD^600^ of 0.2 at 600 nm. The ROS assays were performed on Fries 3 agar plates (16 g/L agar) supplemented with 2 mL of trace elements (167 mg/L LiCl, 227 mg/L CuSO_4_.5H_2_O, 34 mg/L H_2_MoO_4_, 72 mg/L MnCl_2_.4H_2_O, 80 mg/L CoCl_2_.6H_2_O). Drops of 3.5 µL of each tested wheat endophyte were plated on three independent Fries 3 plates amended with 0.5, 1, or 2 mM of H_2_O_2_. Plates containing only Fries 3 medium were used as control. Plates were incubated 23 °C and photographed at 5 dpi. The colony diameter was calculated from digital images of colonies formed in three independent Petri dishes using ImageJ software (47). The percentage of growth inhibition or promotion in response to the presence of H_2_O_2_ was calculated based on the colony diameter of the colonies growing on control plates. These percentages were then plotted in a bubble plot using ggplot2 package from R (50).

The plant secondary metabolite assays were conducted on microtiter plates containing 200 µL of Fries 3 liquid medium supplemented with trace elements and 1 mg/mL at final concentration of CA, or 4 mg/mL at final concentration of NA or QA (all compounds from Sigma, Missouri, USA). The doses were pre-defined in a previous experiments using 10 microbial strains (data not shown). The secondary metabolites CA and NA were solubilized in ethanol, while QA was dissolved in dimethyl sulfoxide (DMSO). Therefore, microtiter wells containing solely Fries 3 medium and Fries 3 supplemented with ethanol or DMSO were used as controls. Each tested wheat endophyte was adjusted to a final OD^600^ of 0.1. Eight and twelve technical replicates were performed for the controls and each secondary metabolites per microbial strains, respectively. Plates were incubated on a rotary shaker (190 rpm) at 23°C and OD^600^ measurements were taken after 24, 48, 72, 96 and 168h of incubation. The growth ratio of each microbial strain was assessed by dividing the mean OD in the presence of each secondary metabolite by the mean OD of the corresponding control culture at each time point. This ratio represents the relative growth of microbial strains in the presence of CA, NA, and QA compared to the control. An OD ratio := 1.1 indicates that the compound promoted microbial growth, while OD ratio :: 0.9 suggests that the compound inhibited the microbial growth. We considered stable bacterial growth if the OD ratio between 0.9 and 1.1. The histogram containing the relative growth ratio distribution was plotted using ggplot2 in R (50). The number of microbial strains showing growth promotion, inhibition, or non-affected growth (ratio between 0.9 and 1.1) in the presence of each secondary metabolite and time point were plotted in an alluvial plot using ggplot2 in R (50).

## Results

### LC-MS/MS analyses reveal qualitative and quantitative effects of *Z. tritici* infection on the wheat apoplastic proteome

To identify the proteins expressed during the compatible and incompatible interactions, we firstly conducted an infection assay with the susceptible wheat cultivar Obelisk (without the resistance gene *Stb6*) and the resistant landrace Chinese Spring (with the resistance gene *Stb6*) inoculated with the IPO323 *Z. tritici* strain, which produce the effector protein AvrStb6 (17, 22, 23, 51). We isolated the apoplastic fluids at 8 dpi, which represents the onset of more extensive fungal intracellular growth, but still at the symptomless biotrophic phase (46, 52). We then confirmed the success of infection at 14 dpi by showing that the IPO323 strain was able to infect and reproduce in leaves of Obelisk, but the infection was aborted in leaves of the resistant Chinese Spring (Fig. S2), corroborating with previous studies (10, 17, 22, 46).

In order to unveil the proteins involved in the wheat immune response against *Z. tritici*, we applied a bottom-up proteomic analysis on the apoplastic fluid extracts (Fig. 1A). A total of 2’455 protein sequences (Fig. S1B, Fig. 1A, and Table S1) were identified, of which 956 (39%) were predicted as apoplastic proteins using SignalP 6.0 and ApoplastP 1.0 (Fig. 1A). Although RuBisCo was not detected in the apoplastic fluids using Western blot analysis (Fig. S1D), it is not surprising that cytosolic proteins such as ribosomal, mitochondrial and chloroplast proteins were identified due to the high sensitivity of the LC-MS/MS approach (Table S2). Some proteins that were not predicted as apoplastic might be secreted by non-conventional secretion mechanisms. Moreover, non-apoplastic proteins may be critical components of plant cells and play a variety of role in plant growth, development, and stress responses (53–55), thus we included these proteins in the further analysis.

**Figure 1.**
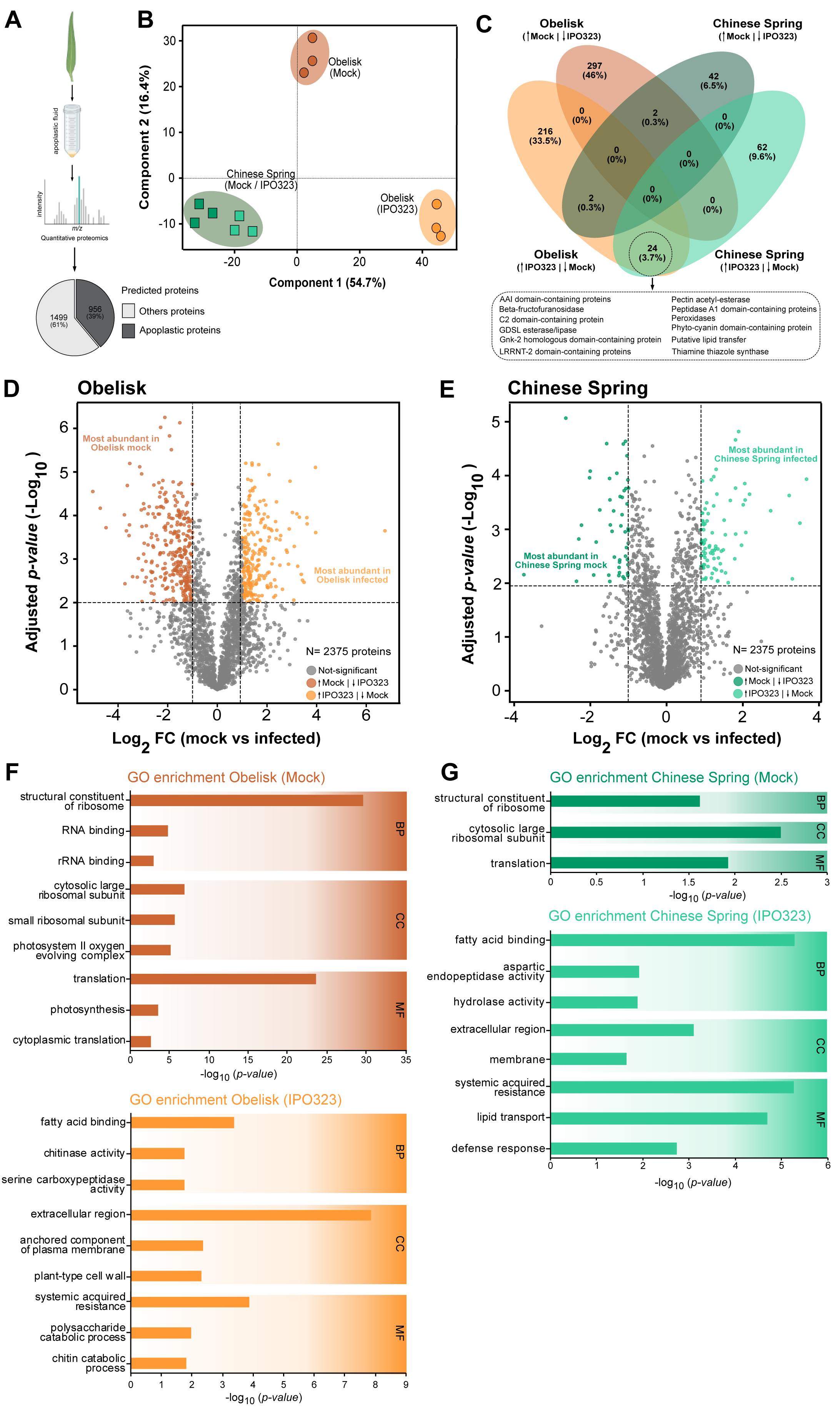
Proteomic analysis to study the impact of the *Zymoseptoria tritici* on the apoplastic proteome of the susceptible and resistant wheat genotypes. (A) Scheme showing the quantitative proteomics used in this study. The pie chart demonstrated the 2’455 total proteins identified in the wheat apoplastic fluids. (B) Principal component analysis (PCA) of apoplastic proteomes of mock-treated and infected Obelisk and Chinese Spring plants. (C) Venn diagrams of 673 differentially expressed proteins in response to *Z. tritici* infection and shared between groups. In highlight, the 24 upregulated proteins exclusively found in Obelisk and Chinese Spring infected plants. (D) Volcano plot of differentially expressed proteins in the susceptible Obelisk. Light orange points represented the 242 proteins that were up-regulated in Obelisk infected plants (adjusted p-value <0.01; FC >2). The dark orange points represented the 299 proteins that were down-regulated in Obelisk during fungal infection (adjusted p-value <0.01; FC >2). (E) Volcano plot of differentially expressed proteins in the susceptible Chinese Spring. Light green points represented the 86 proteins that were up-regulated in Chinese Spring infected plants (adjusted p-value <0.01; FC >2). The dark green points represented the 46 proteins that were down- regulated in Chinese Spring during fungal infection (adjusted p-value <0.01; FC >2). (F) Gene ontology (GO) enrichment terms for down-regulated (dark orange) and up- regulated (light orange) proteins in the susceptible Obelisk plants during fungal infection, respectively. (G) Gene ontology (GO) enrichment terms for down-regulated (dark green) and up-regulated (light green) proteins in the resistant Chinese Spring plants during fungal infection, respectively.

### *Z. tritici* infection causes substantial change in the apoplastic proteome of the susceptible wheat genotype

We compared the proteome data of Obelisk and Chinese Spring to identify the differences between genotypes and between mock-treated and pathogen-infected plants. The wheat genotypes explained the major difference as seen in the PCA plot (Fig. 1B). Intriguingly, when comparing mock-treated and infected plants, we found that fungal infection did not cause a large shift in the Chinese Spring apoplast proteome, but it resulted in a substantial change in the Obelisk cultivar (Fig. 1B).

We further performed pairwise comparisons to identify the differentially abundant proteins in Obelisk and Chinese Spring plants during fungal infection. To this end, we assessed the fold-change of protein accumulation by comparing the mock-treated and IPO323 infected samples from the same wheat genotype. A total of 248 and 87 proteins were higher in abundance in the infected Obelisk and Chinese Spring, respectively (Table S2). On the other hand, 305 and 46 proteins showed a higher abundance in mock treated apoplast of Obelisk and Chinese Spring, respectively (Table S2). Of the measured proteins in Obelisk, about 36% showed a down-regulation in *Z. tritici*-infected samples (242 proteins, Obelisk mock vs infected), while only 13% of proteins in Chinese Spring were down-regulated in the *Z. tritici* infected leaves (86 proteins, Chinese Spring mock vs infected) (Table S2). We hypothesize that these results reflect the extensive physiological and cellular changes which are caused by colonization of *Z. tritici* in the apoplastic space of the susceptible wheat. In Chinese Spring, hyphae of *Z. tritici* do not extent further than the substomatal cavity.

### Proteomic analysis unveils conserved defense-related signatures in *Triticum aestivum* against *Z. tritici* infection

We next aimed to identify common defense proteins associated with *Z. tritici* infection. We compared the proteins in the two wheat genotypes which showed different abundance in the presence of *Z. tritici*, a total of 673 proteins. We identified a set of 24 upregulated proteins shared between infected Obelisk and Chinese Spring plants (Fig. 1C and Table S3). These proteins are predicted to have different biological roles, as cell wall components (*e.g.,* expansins, leucine-rich-repeat extensins, and pectin acetylesterases), in carbohydrate metabolism (*e.g.,* thiamine thiazole synthase and beta-fructofuranosidase), in lipid metabolism (*e.g.,* non-specific lipid transfer proteins (ns-LTPs) and GDSL esterase/lipase), as proteases (*e.g.,* aspartyl protease), and as catalytic enzymes (*e.g.,* peptidases and peroxidases), and as proteins involved in key processes of plant defense responses, such as the cysteine-rich receptor-like kinase (CRK) and leucine-rich repeat (LRR) motif-containing protein. The two identified CRK proteins harbor a Stress-antifungal domain, which is an extracellular domain that consists of the conserved cysteine-rich motif (C-X8-C-X2-C) that possesses antifungal and salt-stress responsive activities (56, 57). Additionally, in plants LRR domains are well-known to be involved in protein-protein interactions, as well as in development and defense processes (58). We conducted a blastp analysis to further predict the functional relevance of the proteins. Hereby we find that the LRR-motif-containing proteins are, polygalacturonase-inhibiting protein (PGIPs) containing LRR-motifs. PGIPs were previously shown to be involved in defense against fungi by inhibiting the activity of polygalacturonases, enzymes secreted by fungal pathogens to degrade pectin in the plant cell wall. Interestingly, pectin degradation by polygalacturonases leads to an accumulation of short-chain oligogalacturonides (OGs) that may be recognized as damage-associated molecular patterns (DAMPs) and elicit plant defense responses (59, 60).

We performed Gene ontology (GO) enrichment analysis on the significantly different proteins (mock vs infected) in Chinese Spring and Obelisk (Fig. 1D-G), and found that six out of seven previously described ns-LPTs (protein IDs A0A3B6AZK3, A0A3B6C6Z8, A0A3B6EM77, A0A3B6RLW5, A0A3B6SLE4, V9I0Q4, Table S3) were annotated to the GO term systemic acquired resistance (SAR) (Tables S4). The role of ns-LTPs in SAR relates to their participation in the intercellular transport of lipids, which serves as crucial signaling molecules in the induction and propagation of defense responses (61). We also pinpoint that the fungus causes a significantly increase in several stress/defense-related proteins in the susceptible Obelisk including endo-chitinases and chitinases (GO cell wall and chitin processes), Bowman-Birk type proteinase inhibitor B5, pectin acetylesterases, Nepenthesin-like aspartic proteinases, and expansins (GO extracellular region), glucan-endo-1,3-beta glucosidase (GO plasma membrane component), peroxidases (GO extracellular region and plant-type cell wall), serine carboxypeptidase (GO serine-type carboxypeptidase activity) (Figs. 1D-F and Table S4). Interestingly, the resistant Chinese Spring also show an increase in the abundance of expansins and peroxidases (GO extracellular region) (Table S4), but the set of proteins is different from Obelisk as we also find an enrichment of proteins of another category of ns-LTPs (GO lipid transport and integral component of membrane), defensins and thaumatin (GO defense response), aspartyl proteases (GO extracellular region and aspartic-type-endopeptidase activity), and GDSL esterase/lipase proteins (GO hydrolase activity) (Figs. 1E-G and Table S4). In summary, our proteome data provide a conserved protein catalog associated with an immune response of *Triticum aestivum* against *Z. tritici* infection, and thereby a novel set of proteins to be functionally validated in future studies.

### Impact of fungal infection on the photosynthesis apparatus and general plant physiological processes

We found that *Z. tritici* infection strongly impacts the abundance of proteins associated with plant physiological processes. First of all, we observed a downregulation of proteins involved in ribosome biogenesis (*e.g.,* 40S and 60S ribosomal proteins) in both Obelisk and Chinese Spring infected plants (Figs. 1E-G and Table 5). The interferences in the translation machinery may be attributed to several direct and indirect consequences of fungal infection, including the activity of fungal effector molecules that may directly target ribosome functions and reduce overall translation (62–64); or it may be attributed to hormone signaling pathways (*e.g.,* salicylic acid and jasmonic acid) involved in defense responses but also affecting ribosomal protein expression (65). Additionally, the downregulation of 40S and 60S ribosomal proteins may be caused by the reallocation of resources involved in the synthesis of specific defense-related proteins and other components necessary for combating the fungal infection (66). We also observed that *Z. tritici* infection interferes with photosynthesis processes in the susceptible Obelisk wheat (Figs. 1D-F and Table S5). During this compatible interaction, we observed the downregulation of the 30S and 50S chloroplast ribosomal proteins, and proteins involved in chloroplast thylakoid luminal, photosynthesis I and II (*e.g.,* PsbP-like proteins) Figs. 1D-F and Table S5. Contrarily, the presence of the *Z. tritici* does not affect the expression of photosynthesis-related proteins in the Chinese Spring (Figs. 1D-F and Table S5). This finding aligns with a previous study demonstrating that the interaction between AvrStb6 and Stb6 proteins in resistant wheat preserves the efficacy of the photosynthesis apparatus (24). Our findings provide novel evidence for the impact of *Z. tritici* on basal physiological processes, beyond defense, in the infected plant tissue during biotrophic colonization of the fungus.

### Apoplast proteome data reveals three novel *Z. tritici* proteins as candidates to fungal biology and pathogenesis regulation

Although several peptides match to protein sequences of the pathogen databases, we consider it more likely that identified peptides have a plant origin. This is due to the fact, that the amount of plant biomass during biotrophic colonization is much higher than the amount of fungal biomass. Despite this, we successfully identified three proteins with peptides exclusively matching the *Z. tritici* database (Table. S6). The absence of N-terminal signal peptides in both proteins (35), may suggest that they either were extracted from *Z. tritici* cells during in the apoplastic fluid or that they are proteins secreted by the fungus by non-conventional secretion mechanisms. The three fungal proteins include a chorismate synthase (Zt09_12_00295) and a cryptochrome/photolyase (Zt09_1_00976) protein, which are both upregulated in Obelisk. The cryptochrome/photolyase protein is also upregulated in Chinese Spring.

The gene encoding the cryptochrome/photolyase protein has previously been identified as a pathogenicity related factor, as it found to be significantly upregulated in an RNA-seq-based transcriptome dataset of IPO323 strain infecting another susceptible wheat cultivar, ‘Riband’ (67). Analysis of the protein sequence by EffectorP 3.0 (68) predicts cryptochrome/photolyase protein to be a cytoplasmic effector further suggesting a relevance of the protein for host infection. The third fungal protein that we identified, belongs to the glycoside hydrolase family 5 (Zt09_11_00165), and was found to be downregulated during fungal infection in Obelisk. Further functional studies are required to elucidate the potential roles of these three novel proteins during the wheat-*Z. tritici* interaction.

### Isolation and characterization of wheat-apoplast microbial communities in Chinese Spring and Obelisk

Above we show that *Z. tritici* infection triggers the secretion of various proteins with antimicrobial properties. These proteins may directly impact the microbial community residing within the wheat apoplastic space. Moreover, the recognition of DAMPs or microbe-associated molecular patterns (MAMPs) within the apoplast compartment can activate pattern-triggered immunity (PTI) and/or effector-triggered immunity (ETI), and these immune responses may also promote the biosynthesis of plant metabolites and compounds possessing antimicrobial activity. Thus, we next hypothesized that these immune-related proteins/compounds could have a direct and/or indirect effect on the plant-associated microbiota leading to the prevalence of specific microbial taxa that tolerate wheat defense responses. To test this hypothesis, we firstly cultured and purified microbial isolates from the apoplastic fluids of mock-treated and IPO323 infected Obelisk and Chinese Spring wheat plants (Fig. S1C-D). In total, we picked 667 isolates cultured from PSM (n= 221, 33.1%), R2A (n= 225, 33.7%), and TSA (n= 221, 33.1%) (Fig. 2A), suggesting a similar cultivation success for the different media. The wheat-enriched endophytic community-based culture collection is composed of 188 purified and archived isolates which we also classified taxonomically using 16S rRNA gene sequencing (Table S7). The 16S rRNA data allowed us to group strains belonging to the same species. Thereby we find that the collection represents 59 species from 21 bacterial families in the phyla Actinobacteria (n= 42; 13 species), Bacteriodetes (n= 8; 6 species), Firmicutes (n= 1; 1 specie), and Proteobacteria (n= 124; 36 species). We also isolated fungi including Ascomycetes (n= 9; representing the re-isolation of *Z. tritici* from Obelisk and Chinese Spring infected plants) and Basidiomycetes (n= 4; 3 species) (Figs. 2A-B and D, and Table S7). Importantly, *Z. tritici* was exclusively re-isolated from infected Obelisk and Chinese Spring plants, confirming i) the presence of *Z. tritici* in the apoplast of infected plants and ii) the absence of cross-contamination between mock and inoculated plants. Overall, the predominant phyla in the culture collection were Proteobacteria (65.9%), followed by Actinobacteria (22.3%), Ascomycetes (4.7%), Bacteroidetes (4.2%), Basidiomycetes (2.1%), and Firmicutes (0.5%).

**Figure 2.**
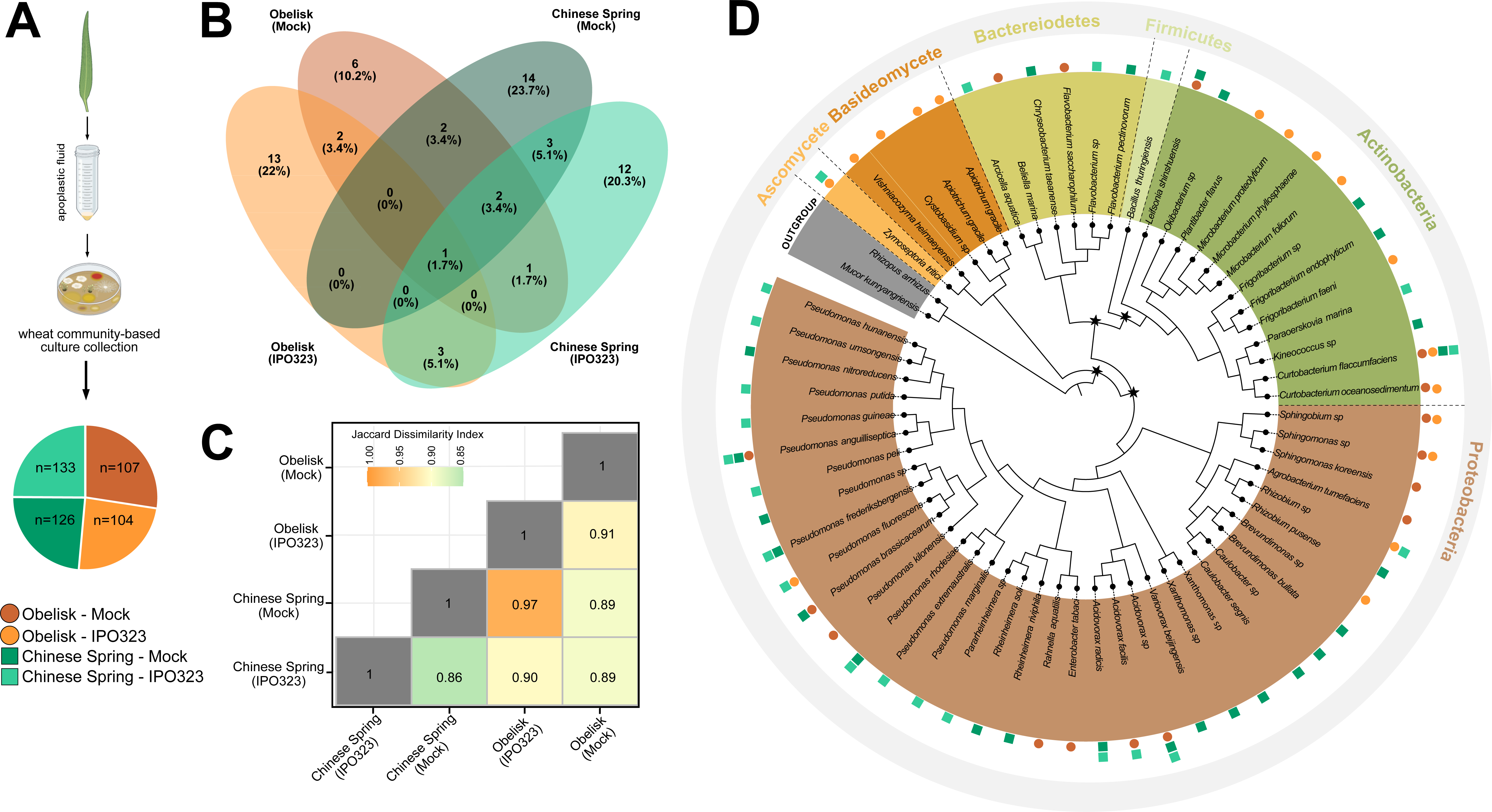
Wheat-enriched endophytic community-based culture. (A) Scheme showing the microbial isolation procedure used in this study. The pie chart demonstrated the 667 microbes isolated from the wheat apoplastic fluids, and the respective number of isolates per groups (mock-treated or infected Obelisk and Chinese Spring). (B) Wheat-enriched endophytic community-based culture collection is composed of 188 purified and archived isolates which were classified taxonomically using partial 16S rRNA and ITS sequences. These 188 microbes represent 59 species. Venn diagrams showing the number of shared OTUs between groups. (C) Heatmap showing Jaccard index to compare the dissimilarity between microbial communities based on their species richness. (D) A reference phylogenetic tress of 59 microbial species used in this study. Bar colors represent the corresponding phyla in all figures in this study. Black star indicates the ancestral of each phylum. Circles and squares represent the wheat genotypes and treatment from where the microbes were isolated. Dark or light orange circles indicated the microbe isolated from mock-treated or *Z. tritic*i infected Obelisk plants, respectively. Dark or light green squares indicated the microbe isolated from mock-treated or *Z. tritic*i infected Chinese Spring plants, respectively.

### *Z. tritici* infection does not affected microbial species richness in resistant host, but leads to more stochastic microbial communities in a susceptible host

A previous study showed that IPO323 infection promoted a significant reduction in community richness in Chinese Spring, but the fungal infection had no significant effect on community composition of the susceptible cultivar Obelisk (10). With our culture-based approach we obtain a similar number of OTUs (23) from mock- and *Z. tritici* infected Chinese Spring plants. For Obelisk we isolated 14 and 17 OTUs from mock and *Z. tritici* infected plants, respectively. Although, the culture-based approach does not provide the complete diversity of apoplast-associated microbes, we considered the culture collections as proxies for the complete diversity. Hereby, we compared the composition of microbes isolated from the different conditions and used the Jaccard index (JI) to compare the dissimilarity between microbial communities based on their species richness (Fig. 2C). Comparing the effects of *Z. tritici* infection in the two wheat genotypes, we find the highest dissimilarity between Chinese Spring-mock and Chinese Spring-infected microbial communities (JI=0.86) The dissimilarity between Obelisk mock and infected communities was considerably lower (JI=0.91). This observation is in agreement with our previous study which likewise showed the strongest impact of *Z. tritici* infection in the resistant wheat Chinese Spring (10). This observation correlated with an excess of defense related metabolites in pathogen infected leaves.

### Identification of core microbes which show increased resilience to wheat defense responses

The microbes recovered from mock-treated Obelisk plants showed a predominance of Proteobacteria (70.3%), followed by Actinobacteria (24.3%) and Bacteroidetes (5.4%). However, a distinct shift in microbial composition was observed in infected Obelisk plants, with Actinobacteria dominating 60.4%, while Proteobacteria accounted for only 11.6%. Actinobacterial species have been extensively studied for their plant-beneficial roles in disease suppression and as keystone taxa in plant microbial communities (69–72). Within our culture collection, all Actinobacterial representatives belonged to the *Microbacteriaceae* family; however, we observed variations at the genus level between mock-treated and infected Obelisk plants. While *Curtobacterium* and *Leifsonia* were identified in mock-treated plants, infected Obelisk plants were colonized by the genera *Curtobacterium*, *Kineococcus*, *Frigoribacterium*, *Microbacterium*, and *Plantibacter*. Interesting, when comparing the associated microbes of Obelisk, we found that *Curtobacterium* and *Sphingomonas* were the only genera recovered from both uninfected and infected (Fig. 2B), suggesting their resilience to the plant defense responses triggered by *Z. tritici* infection in this susceptible wheat cultivar or the antimicrobial compounds produced by the fungus itself.

We next evaluated the recovered microbial communities associated with Chinese Spring (Fig. S3 and Table S7). Proteobacteria was the most predominant phylum in both mock-treated (83.7%) and infected (81.3%) plants, followed by Actinobacteria (8.1% and 11.8%, respectively) and Bacteroidetes (8.1% and 3.4%, respectively). Additionally, the phylum Firmicutes (1.7%) was identified solely in Chinese Spring infected plants. When comparing Proteobacteria at the genus level, *Pseudomonas* was the predominant genus in both mock-treated and infected plants (60.9% and 79.1%, respectively). As a point of comparison, *Pseudomonas* species accounted for 53.8% of the recovered microbiota from mock-treated Obelisk plants compared to only 20% in this susceptible cultivar infected with *Z. tritici*. Finally, by comparing the associated microbes of Chinese Spring, we found that *Curtobacterium*, *Acidovorax*, and *Pseudomonas* were the three genera recovered from both uninfected and infected plants (Fig. 2B), indicating their resilience to the plant defense responses triggered by *Z. tritici* infection in this resistant wheat genotype or the antimicrobial compounds produced by the fungus itself. In summary, *Z. tritici* infection in the susceptible host resulted in a shift towards Actinobacteria dominance and a decrease in Proteobacteria members, while *Pseudomonas* predominated in other host conditions. These findings highlight the interplay between host susceptibility and microbial community composition, underscoring the potential importance of Actinobacteria and *Pseudomonas* in plant defense responses and disease outcomes.

### Microbial *in-vitro* confrontation assays identify antagonists of *Z. tritici* in the wheat apoplast

We next sought to characterize the interaction response of recovered microbes and *Z. tritici* in an in vitro assay. Briefly, we conducted confrontation co-cultures to explore the antagonistic potential of wheat endophytic microbes against three geographically distinct *Z. tritici* isolates, including the Dutch isolate IPO323, the Danish Zt05, and the Iranian Zt10. Scheme illustrating the plates is shown in Fig. 3A, and the overall inhibitory phenotypes summarized in Figs. S4-6.

**Figure 3.**
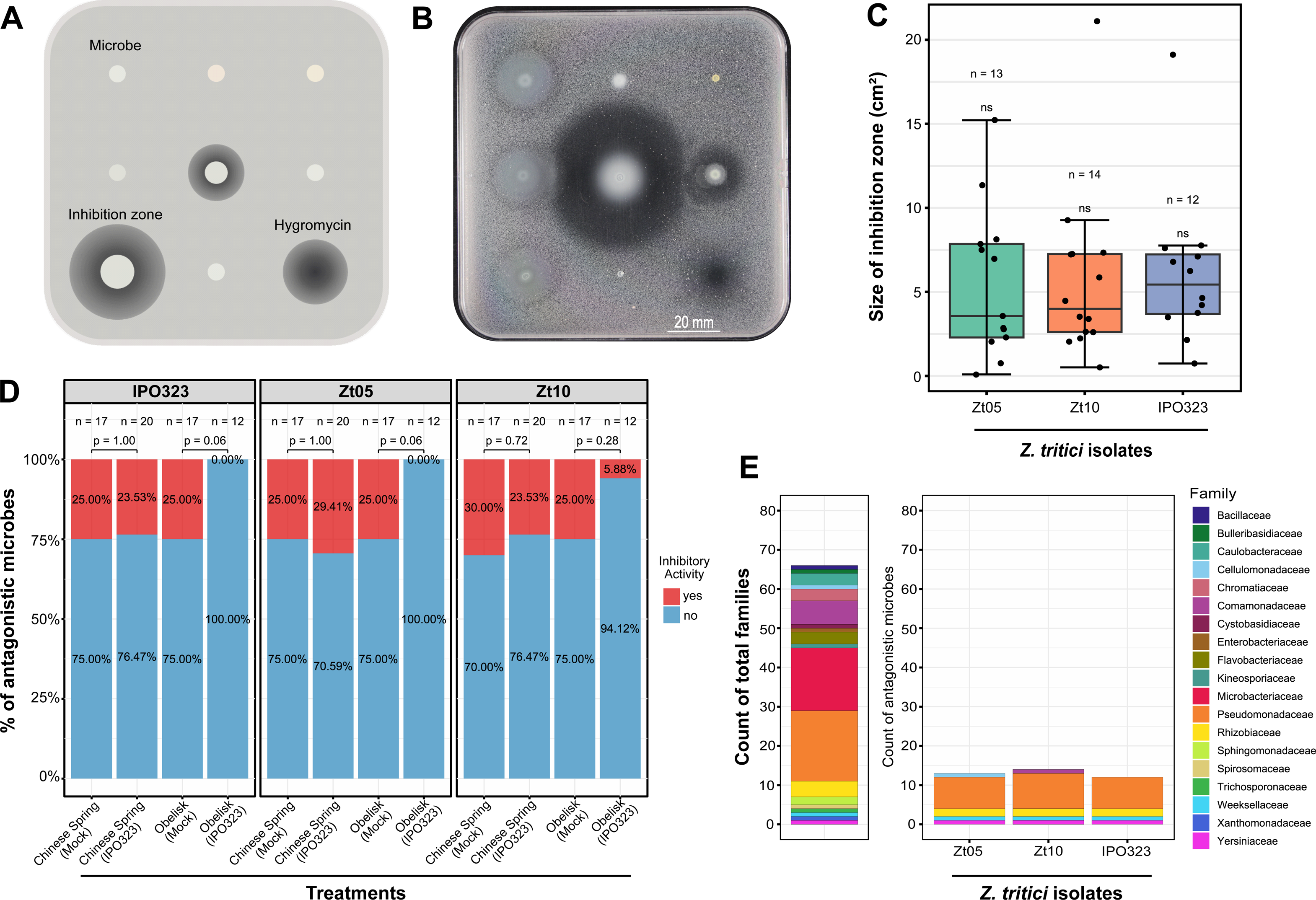
The apoplast of wheat plants harbor an arsenal of potential antagonistic endophytes against *Zymoseptoria tritici*. (A-B) Scheme and exemplification of the antagonistic assay performed *in vitro.* Plates containing solely Fries 3 medium were used as control to ensure microbial growth. Hygromycin B was used as positive control for fungal growth. (C) Three *Z. tritici* isolates were used in this study, such as the Dutch isolate IPO323, the Danish Zt05, and the Iranian Zt10. Roughly 20% of the wheat-recovered microbes were able to inhibited at least one of the *Z. tritici* isolates. Black points represent antagonistic microbes. Isolated that did not inhibited *Z. tritici* were excluded. The size of inhibition zones did not differ among the three tested *Z. tritici* isolates based on the Kruskal-Wallis statical analysis. (D) Proportion of microbes isolated from different wheat genotypes varying in the inhibitory activity against *Z. tritici*. (E) Composition of the wheat-enriched endophytic community- based culture (left) and the microbes inhibiting the three different *Z. tritici* isolates (right). Microbes were colored based on their respective family classification.

Six wheat-associated microbes did not show visible growth (*e.g.,* Fig. 3B), and were therefore excluded from the analysis. In general, we did not observe any substantial difference in the inhibitory activity among fungal isolates suggesting that sensitivity and tolerance to plant-associated bacteria overall are conserved mechanisms in *Z. tritici*, at least under the conditions tested here (Fig. 3C and S7A). The assay revealed an inhibitory effect in 15 out of 72 microbes against at least one of the tested *Z. tritici* isolates. Twelve isolated inhibited all the three *Z. tritici* isolates, while the *Paraoerskovia marina* affected negatively the growth of Zt05, and *Variovorax beijingensis* and *Pseudomonas fluorescens* inhibited growth exclusively for Zt10. Although the size of inhibition zones greatly ranged from 21 cm² for *Pseudomonas brassicacearum* to 0.09 cm² for *Paraoerskovia marina*, we found no statistical differences for the size of inhibition zones among the three *Z. tritici* isolates for the individual bacterial associations (Fig. 3C and S7A).

We further asked whether the antagonistic interactions with *Z. tritici* included specific microbial families. Among the 19 microbial families representing the members from wheat-associated microbes, six families were capable of inhibiting at least one of the *Z. tritici* strains, such as *Comamonadaceae*, *Cellulomonadaceae*, *Pseudomonadaceae*, *Rhizobiaceae*, *Weeksellaceae*, and *Yersiniaceae*, the *Pseudomonadaceae* being the most prevalent inhibitory bacterial family. Other less represented families, such as *Yersiniaceae* and *Weeksellaceae*, also caused strong *Z. tritici* growth inhibition (Fig. 3E). We also find that a significant majority of the microbes inhibiting the three tested *Z. tritici* isolates are gram-negative bacteria (Fig. S7B). Interestingly, we noticed a pattern in the proportion of antagonistic microbes based on the wheat genotype and infection condition from which they were isolated. Microbes isolated from the infected susceptible Obelisk plants were the lesser inhibitors of *Z. tritici* compared to the other treatments (Fig. 3D). In fact, only *Pseudomonas fluorescens* inhibited Zt10, while none of the microbes isolated from *Z. tritici*-infected Obelisk inhibited Zt05 or IPO323. Contrarily, the significant proportions of antagonistic microbes were isolated from both mock-treated Obelisk and Chinese Spring, and from the incompatible interaction (Fig. 3D). Here, we identified wheat endophytic microbes with antagonistic potential against different *Z. tritici* isolates, suggesting the necessity of further investigation to elucidate their specific mechanisms in controlling fungal growth.

### Effect of defense-related plant compounds on apoplast-associated microbes

The defense response of plants to biotic stresses includes the production of ROS and defense-related secondary metabolites. While triggered by pathogenic microbes, the plant defense response may also impact the diversity of commensal microbes colonizing plant tissues (73). To investigate the specific adaptations of the apoplast microbes isolated from Obelisk and Chinese Spring, we phenotyped 69 microbes, including the pathogen *Z. tritici*, for their tolerance or susceptibility to four representative exogenous sources of plant defense compounds (H_2_O_2_, CA, QA, and NA). The phenotyping consisted of growth measures (i) on solid medium at a single timepoint and (ii) in microtiter plates over a time series.

In the first assay based on microbial growth on solid medium amended with H_2_O_2_, we found a significant variation in growth across different bacterial genera and species (Fig. 4). For example, all members from the genera *Acidovorax* and *Brevundimonas* were fully inhibited at 0.5 µM of H_2_O_2_, while bacteria belonging to the genera *Rahnella* and *Enterobacter* only were inhibited at 1 µM. All tested microbes were inhibited at 2 µM, except for *Z. tritici* and one *Pseudomonas anguilliseptica* (AF0034) isolated from mock-treated Obelisk plants. Genes associated with increased tolerance to H_2_O_2_ were previously identified in *Z. tritici* based on a GWAS study (74); but the ROS-scavenging mechanism for *P. anguilliseptica* remains unknown. Furthermore, we also observed within species variation in the tolerance levels to H_2_O_2_ including a comparison of different strains of *Pseudomonas spp.* and *Flavobacterium spp.* (Fig. 4). While the growth of *Pseudomonas anguilliseptica* or *Flavobacterium hydatis* were promoted at 0.5 and 1 µM, the *Pseudomonas brassicacearum* and *Flavobacterium pectinovorum* were inhibited at 0.5 µM (Fig. 4). Interestingly, strains of Actinobacteria showed a higher tolerance to H_2_O_2_ by showing either a slight growth promotion or a slight inhibition compared to the rest of the wheat-associated microbes Fig. 4). In our study, the majority of Actinobacterial strains were obtained from Z. tritici-infected Obelisk leaves (Fig. S3B). This observation indicates a more efficient, and potential adaptive, ROS-scavenging mechanism in Actinobacteria allowing these bacteria to cope with the oxidative stress induced during *Z. tritici* infection. Finally, the fungal isolates also displayed high tolerance to H_2_O_2._ Our assay included fungal species belonging to the genera *Apiotrichum*, *Cystobasidium*, and *Vishniacozyma*. The enhanced ability of fungi to tolerate oxidative stress is known to be attributed to their intricate antioxidant machinery and repair mechanisms, enabling them to survive in environments with high oxidative potential (74–77)

**Figure 4.**
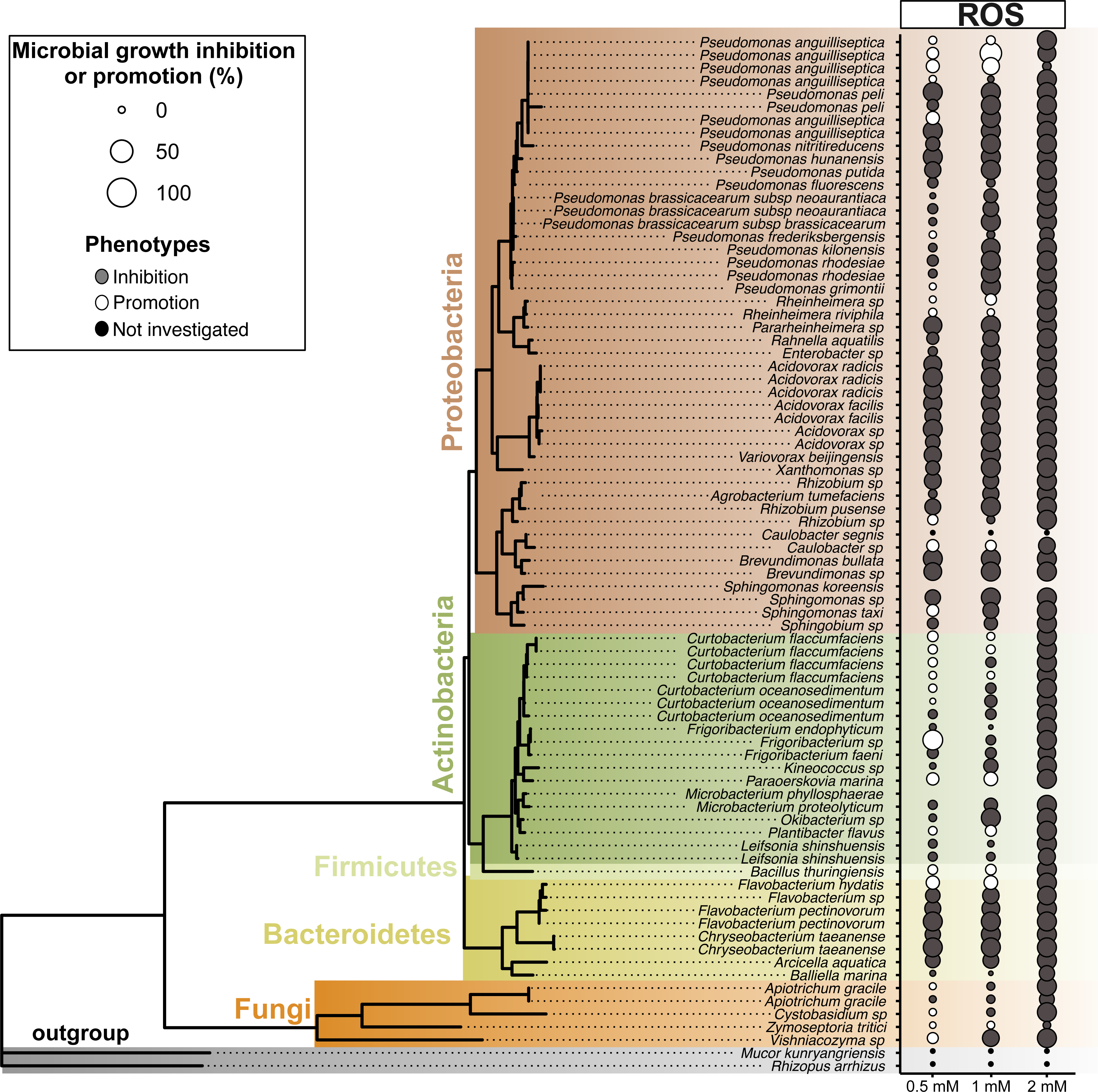
Phylogenetic tree depicting the evolutionary relationship of all retrieved microbial species and assessment of reactive oxygen species (ROS) sensitivity. The phylogenetic tree was reconstructed based on partial 16S rRNA and ITS sequences. Nodes are supported based on 500 bootstraps performed in RAxML. Two fungal species *Rhizopus arrhizus* and *Mucor kunryamgriensis* from the Mucoroycota were used as outgroups in the tree. Three doses of hydrogen peroxide (H_2_O_2_) were used in solid growth media for assessing the growth sensitivity of 69 endophytic strains retrieved from wheat. The percentage of growth inhibition and promotion is based on the ratio between colony diameter under H_2_O_2_ and colony diameter of colonies growing on control plates. Phenotyping performed at 5 days post inoculation.

In the second assay, we determined microbial growth based on estimates from microtiter plates. Hereby, we assessed the impact of CA, QA, and NA on microbial growth. Also, we measured growth promotion, growth inhibition and no-effect on growth across all 69 apoplast-associated microbes over five timepoints (24, 48, 72, 96 and 168 hours). Considering the entire course of the experiment, each of the 69 microbes were evaluated five times, resulting in 345 measurements. We initially noted a variation in the frequency of microbial isolates that showed inhibition, promotion and stable (no change compared to control) phenotypes across compounds and timepoints (Fig. 5 and Table S8).

**Figure 5.**
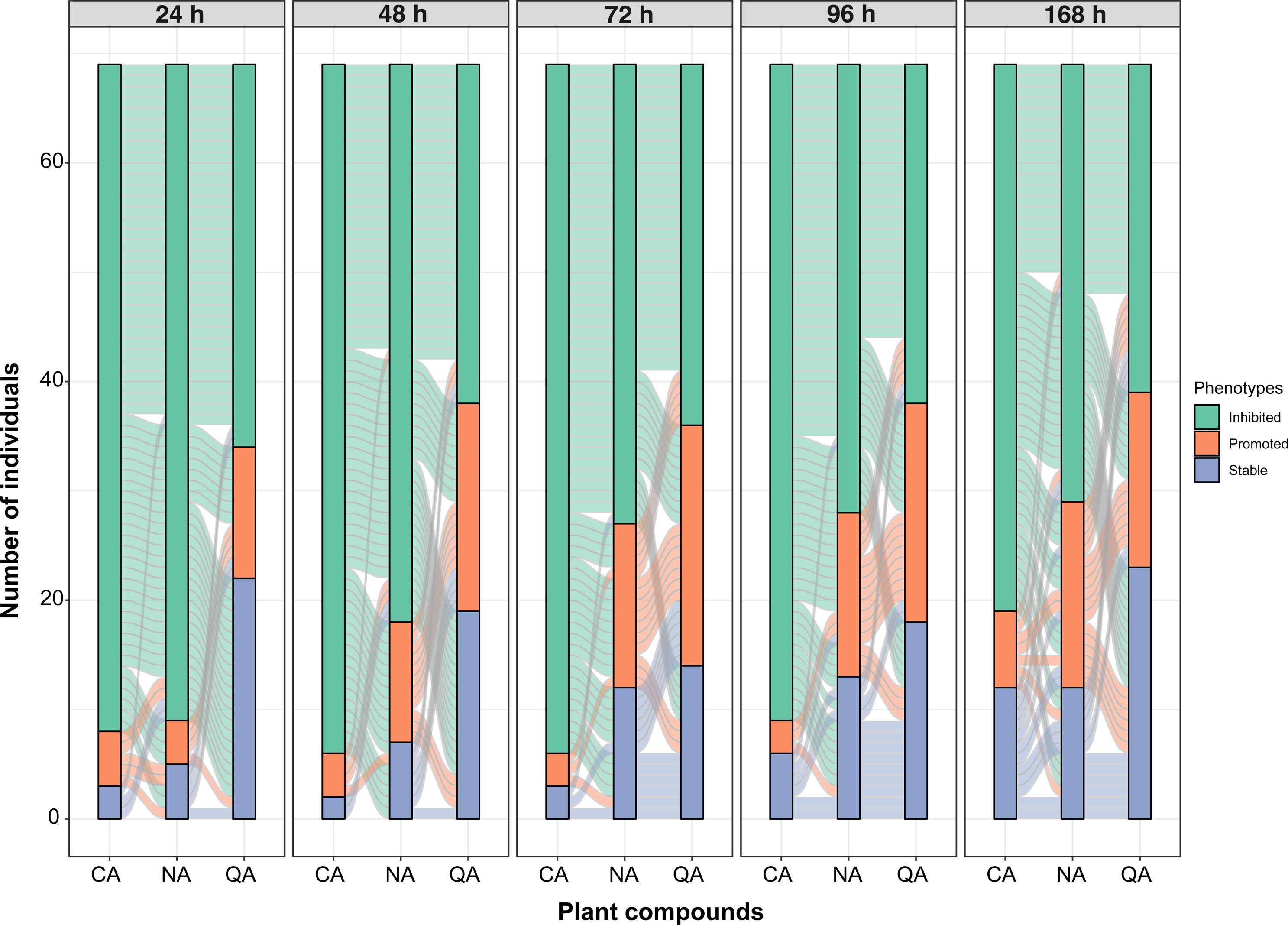
Phenotyping of 69 strains for growth inhibition, promotion, and stability across five time points and three tested exogenous plant secondary metabolites. A total of 69 isolates representing 59 species were tested in this study. The three used plant secondary metabolites are cinnamyl alcohol (CA), quinic acid (QA), and nonanoic acid (NA). The five time points are 24, 48, 72, 96 and 168 hours after inoculation of microtiter plates. In the alluvial plot, the three strata represent each phenotype per plant metabolites within a time point, and the connections illustrate the phenotype of a given microbe between strata. Phenotypes were defined by dividing the OD of a given strain growing under a plant metabolite by its OD when growing in the control media. An OD ratio := 1.1 indicates growth promotion, :: 0.9 growth inhibition and in between stable growth.

Overall, the compound CA exhibited the highest levels of inhibition with 86% of the measurements showing a growth inhibition (297 inhibitions out of 345), followed by NOA with 67,8% of measurements (234 inhibitions out of 345), and QA with the lowest levels of inhibition, 46,3% (160 inhibitions out of 345) (Fig. 5 and Table S8). These observations were consistent across all time points with the compound CA displaying the highest levels of inhibition (Fig. 5 and Table S8). Interestingly, we highlight intrinsic differences of individual microbes to the three tested compounds, whereas a given individual may show inhibition or promotion of growth according to different compounds.

We next investigated in more detail how individual microbes varied in susceptibility to each compound overtime (from 24 to 168 hours post incubation). For CA, *Pseudomonas kilonensis* (AF0006), *Enterobacter sp* (AF0002), and *Curtobacterium oceanosedimentum* (AF0066) exhibited more growth promotion than any other strain, while *Sphingobium bisphenolicum* (AF0048), *Agrobacterium tumefaciens* (AF0129), *Acidovorax sp* (AF0089), *Pseudomonas sp* (AF0130), *Brevundimonas sp* (AF0072), and *Pseudomonas brassicacearum* (AF0120) were the most inhibited strains (Table S9). Remarkably, none of those abovementioned strains were isolated from Chinese Spring infected plants.

For NA, *Pararheinheimera sp* (AF0141), *Pseudomonas hunanensis* (AF0142), *Baliella marina* (AF0012*), Flavobacterium hydatis* (AF0027), *Microbacterium sp* (AF0077), *Bacillus thuringiensis* (AF0154), and *Caulobacter sp* (AF0118) were the most promoted strains, while *Acidovorax sp* (AF0089), *Curtobacterium flaccumfaciens* (AF0165), *Variovorax beijingensis* (AF0128), *Curtobacterium oceanosedimentum* (AF0004), *Kineococus sp* (AF0060), *Pseudomonas anguilliseptica* (AF0162), *Rhizobium sp* (AF0164), and *Leifsonia shinshuensis* (AF0088) were the most inhibited strains (Table S9). The microbes AF0141, AF0142, AF0154, AF0162, AF0164, and AF0165 were isolated from *Z. tritici*-infected Chinese Spring plants, and we hypothesized that *Pararheinheimera sp, Pseudomonas hunanensis,* and *Bacillus thuringiensis* may benefit from the accumulation of nonanoic acid in the apoplast and these microbes may potentially play a role in the plant-pathogen interaction. On the other hand, *Pseudomonas anguilliseptica, Rhizobium sp*, and *Curtobacterium flaccumfaciens* may be sensitive to the pathogen-induced immune response.

Finally, for QA, we observed that *Pseudomonas putida* (AF0171), *Pseudomonas hunanensis* (AF0142), *Pararheinheimera sp* (AF0141), *Pseudomonas umsongensis* (AF0144), *Chrysepbacterium taeanense* (AF0131), *Pseudomonas anguilliseptica* (AF0138), *Acidovorax facilis* (AF0014), and *Pseudomonas rhodesiae* (AF0111 and AF0139) were the most promoted strains, while *Pseudomonas sp* (AF0130), *Agrobacterium tumefaciens* (AF0129), *Pseudomonas frederiksbergensis* (AF0191), *Pseudomonas brassicacearum* (AF0029, AF0102, and AF0120), *Sphingobium bisphenolicum* (AF0048), and *Microbacterium proteolyticum* (AF0059) were the most inhibited strains (Table S9). The microbes AF0171, AF0138, AF0139, AF0141, AF0142, AF0144, and AF0191 were isolated from *Z. tritici*-infected Chinese Spring plants, and we hypothesized that different *Pseudomonas* and *Pararheinheimera* species may benefit from the production of quinic acid during the induced defense response. Contrarily, *Pseudomonas frederiksbergensis* appears to be sensitive to the defense related compound.

In summary, *Pseudomonas hunanensis* (AF0141) and *Pararheinheimera sp* (AF0142) were the two bacteria isolated from *Z. tritici*-infected Chinese Spring plants which showed growth promotion when exposed to both NA and QA, while *Agrobacterium tumefaciens* (AF0129), *Acidovorax sp* (AF0089), and *Pseudomonas brassicacearum* (AF0120) isolated from the mock-treated Chinese Spring plants were the most inhibited microbes (Table S9). The 10 most promoted and inhibited microbes over the time course of the experiment are summarized in Table S9. These findings demonstrate a remarkable variation in the tolerance of apoplast-associated microbes to plant compounds previously shown to be induced during *Z. tritici* infection. Future studies should address the molecular mechanism underlying the growth inhibition and promotion and their impact for plant health and pathogenesis.

## DISCUSSSION

In their natural environment, plants are constantly challenged by pathogenic organisms which attempt to invade below and above ground tissues. Pathogenic organisms can induce immune responses conferring significant alterations in the composition of molecules which are secreted into the apoplastic space. In response, plants produce and secrete a variety of antimicrobial proteins and metabolites to hamper pathogen invasion. How do commensal microorganisms that inhabit the apoplastic space cope with recurrent biotic stress responses? We addressed this question in our study by characterizing fluctuations in biotic conditions in wheat apoplast and by investigating specific adaptations of apoplast-associated microbes.

We successfully isolated apoplastic fluids from the two wheat genotypes, Obelisk and Chinese Spring, allowing us to characterize the biochemical environment between cells in the leaf mesophyll. The two wheat genotypes are genetically different; Obelisk is an inbred cultivar while Chinese Spring is a landrace known to have a broader resistance spectrum towards pathogens (78). In our comparative proteome analysis, first of all, we observed a considerable differentiation in protein composition between the two wheat genotypes. The different genetic-make up of these two wheat genotypes may to some extent explain differences in proteome content; however, we cannot exclude that the two genotypes also differ in the amenability of protein extraction.

Obelisk and Chinese Spring differ in their resistance towards the *Z. tritici* isolate IPO323. This is due to the production of the cell wall associated immune receptor Stb6 in Chinese Spring; a receptor that is activated by the secreted protein AvrStb6 that is produced by IPO323. We previously showed that the abundance of immune-related metabolites is significantly up-regulated in Chinese Spring during early invasion of *Z. tritici* (10). In contrast, the production of immune-related metabolites is suppressed in Obelisk during IPO323 invasion. Proteome analyses of infected Obelisk and Chinese Spring leaves showed a common response to fungal invasion, suggesting a conserved initial response to fungal invasion (Fig. 1C). The ns-LPTs (containing bifunctional inhibitor/plant lipid transfer protein/seed storage helical domain), for example, have been implicated in the regulation of plant defense responses against various pathogens (61). Additionally, peroxidases are upregulated in both wheat genotypes during *Z. tritici* infection. Peroxidases are enzymes are known to play a crucial role in the generation of reactive oxygen species, an initial defense response conferring downstream immune signaling and local plant cell death (79, 80). Although Obelisk is susceptible to the invasion of the *Z. tritici* isolate IPO323, we recognize several defense related responses reflected in an upregulation of endochitinases, chitinases, and a serine carboxypeptidase, each consistent with the activation of defenses against fungal pathogens (81–83). Endochitinases and chitinases are enzymes involved in the degradation of chitin, a component of fungal cell walls, while serine carboxypeptidases have been implicated in the defense against fungal pathogens by cleavage off essential amino acids from fungal cell walls. In contrast, proteins that are specifically upregulated in Chinese Spring reflect a different mechanism of defense responses (*e.g.,* defensins, thaumatin, aspartyl proteases, and GDSL esterase/lipase). Defensins are small cysteine-rich peptides with antifungal activity, while thaumatin is known to have antifungal and antibacterial properties (84–86). Aspartyl proteases play a role in plant cell death and the degradation of invading pathogens, while GDSL esterase/lipases are involved in the regulation of plant defense responses against various pathogens (87, 88). We also observe an upregulation of ns-LTPs in Chinese Spring, which may be an important component of the systemic acquired resistance (SAR) induced by *Z. tritici* in the resistant cultivar (10). In *Arabidopsis*, the ns-LTPs are induced by immune signaling and secreted into the apoplast where they participate in the intercellular, long-distance transport of lipids, acting as crucial signaling molecules in the induction and propagation of defense responses (89). This systemic lipid transfer is believed to prime uninfected tissues, enabling a robust defense response upon subsequent pathogen attack. Furthermore, ns-LTPs have been implicated in enhancing the stability of lipid rafts in the plasma membrane, which act as signaling platforms for immune receptors and downstream defense-related signaling molecules (89, 90). In summary, the proteome data presented here provides an accurate and novel catalogue of conserved and genotype specific apoplast proteins, including shared and specific defense-related proteins. We specifically identify components of systemic immune signaling in Chinese Spring, which may be considered in future breeding strategies. Interestingly, *Z. tritici* appear to be tolerant to a variety of defense related proteins which are upregulated in the susceptible Obelisk, but without effect on pathogen propagation.

From the apoplasic fluid samples we also isolated endophytic bacteria and fungi. Such broader approach allowed us to characterize not only microbial diversity, but also the footprints of microbial adaptations to varying wheat defense responses. Importantly, our studied was designed in a way that the two wheat genotypes were grown in the same soil and thereby provided with the same source of microbial communities. We recognize that the culture-based approach allows us to characterize only a subset of the complete microbiota diversity. Thereby, we can neither make conclusive statements about differences between wheat genotypes or conditions as the sampling efficacy may be biased by our ability to isolate and cultivate specific microbes. In total we isolate 667 microbes comprising 59 different taxa. In our study, we observed that the wheat-recovered microbes were predominantly composed of Gammaproteobacteria within the Proteobacteria phylum. This result is consistent with previous studies reporting the enrichment of Gammaproteobacterial class in various wheat genotypes and plant compartments (91–93). However, we identified a clear and significant shift from Proteobacteria to Actinobacteria specifically during the compatible interaction, where Obelisk wheat was infected by *Zymoseptoria tritici*. This shift suggests that the defense responses elicited by Obelisk may have a detrimental effect on Proteobacteria endophytes. The transition from Proteobacteria to Actinobacteria can have profound implications for the ecosystem’s functioning and plant health. The capacity of Actinobacteria to degrade complex organic compounds, such as cellulose and lignin, can influence nutrient cycling and organic matter decomposition, thereby impacting ecosystem processes. Proteobacteria, particularly Alpha- and Gamma-Proteobacteria, contribute to plant growth promotion, nitrogen fixation, and biocontrol activities, while Actinobacteria are known to assist in plant defense against pathogens (94, 95). Consequently, a shift from Proteobacteria to Actinobacteria may disrupt these beneficial interactions, potentially compromising plant growth, disease resistance, and overall plant health. Furthermore, this shift could lead to reduced functional redundancy and decreased resilience within the microbial community, rendering it more susceptible to environmental disturbances. Additionally, alterations in microbial interactions can occur, influencing the dynamics of the entire microbial community. To comprehensively understand the consequences of the Proteobacteria to Actinobacteria shift, context-specific studies are needed to unravel its effects on ecosystem processes and plant-microbe interactions.

In terms of microbiota disappearance and resilience, we unveiled that *Pseudomonas* were eliminated during the compatible interaction, while *Curtobacterium* and *Sphingomonas* were the only two genera resilient to defense responses of the susceptible Obelisk. In fact, we observed that *Curtobacterium* and *Sphingomonas* species were highly tolerant when challenged against exogenous H_2_O_2_ in our *in vitro* assays. These findings are in agreement with a previous study that demonstrated extensive ROS production *in planta* in a susceptible interaction (96). ROS, generated by plants as defense mechanisms against pathogens and environmental stresses, can significantly influence the colonization and behavior of endophytes. Elevated ROS levels may impede endophytic colonization or induce alterations in microbial gene expression and metabolic pathways. Nevertheless, certain endophytic microbes have evolved mechanisms to tolerate or exploit ROS as signaling molecules (73). Thus, we postulated that the Obelisk-infected associated microbes were more resilient to H_2_O_2_ due to their previous adaptation to cope with oxidative stress generated during wheat development and/or abiotic/biotic stress. Regarding *Pseudomonas*, we observed *in vitro* that this bacterial genus, overall, show a lower tolerance to exogenous H_2_O_2_.

*Pseudomonas* was also the dominant genus recovered from the incompatible interaction. A previous study showed that in an incompatible interaction of Stb6- AvrStb6 free radicals such H_2_O_2_ are neutralized at the biotrophic phase, preventing the oxidative burst (24). Therefore, we hypothesized that the lower concentration of apoplastic H_2_O_2_ in the incompatible interaction enabled *Pseudomonas* species to colonize the apoplastic space of the resistant Chinese Spring plants. However, when evaluating *Pseudomonas* species against three exogenous wheat secondary metabolites, we observed that this bacterial genus displayed the highest tolerance levels among all microbial species. These three secondary metabolites were previously found to be enriched *in planta* during the wheat incompatible interaction (10). Plant-produced secondary metabolites, including phytochemicals and antimicrobial compounds, can impact endophytic microbial communities by either inhibiting or stimulating their growth and activity. We then hypothesized that the predominance of *Pseudomonas* genus in Chinese Spring plants is due to the reduced H_2_O_2_ and higher tolerance to secondary metabolites in the apoplast space. These reciprocal interactions among ROS, secondary metabolites, and endophytic microbes constitute a complex and dynamic relationship that profoundly shapes the composition and functionality of the endophytic microbiome. Another evidence of this complexity is reflected in the high potential of *Pseudomonas* species to inhibit *Z. tritici* growth *in vitro*, while this bacterial genus is depleted in the compatible interaction.

Our findings provide novel evidence for the impact of *Z. tritici* on defense responses and physiological processes in the infected plant tissue. We provided evidences that specific components of plant-immune responses shape microbial profiles leading to the prevalence of microbes adapted to oxidative stress or secondary metabolites, depending on the outcome of the host-pathogen interaction responses. Further studies will be necessary to address the impact of the apparent shift in dominance from Proteobacteria to Actinobacteria in the susceptible Obelisk, as well as the direct endophytic-triggered inhibition of *Z. tritici* growth during plant colonization and the onset of disease.

## Data availability

Proteome data is made available upon request.

Wheat-associated microbes are stored at -80C in the lab of EHS, CAU Kiel and are available upon request.

## Author contributions

Conceptualization: CSF, EHS

Methodology: CSF, MA, LC, MH

Formal analyses and investigation: CSF, MA, CI, and JG

Resources: EHS and AT

Writing of original draft: CSF, EHS

Review and editing: All authors

Funding acquisition: EHS, AT

## Acknowledgements

This research was funded by the German Research Foundation in the framework of the collaborative research center “Origin and Function of Metaorganisms”, CRC1182, sub-project A3 and Z3. We thank Dr. Danilo A. dos Santos Pereira and Dr. Marco Alexandre Guerreiro for sharing R scripts. We also thank Susanne Braun for the remarkable technical support.

## Supplemental Material

**Figure S1. Validation of apoplastic fluids to isolate proteins and microbes associated with the apoplast compartment of wheat plants.** (A) Scheme showing the methodology used for isolation of apoplastic fluids from mock-treated and infected Obelisk and Chinese Spring plants. Apoplastic fluids were then used for microbial isolation and proteomic analysis. (B) A total of 2’455 proteins were identified in the wheat apoplastic fluids. Bar plots illustrate the total proteins isolated from the three biological and two technical replicated for each wheat genotype and treatment. Dark and light orange bars indicate proteins isolated from mock-treated and infected Obelisk plants, respectively. Dark and light green bars indicate the proteins isolated from mock-treated and infected Chinese Spring plants, respectively. (C) A total of 667 microbes were isolated from the apoplastic fluids. Bar plots illustrate the microbes isolated from each wheat genotype and treatment. Dark and light orange bars indicate microbes isolated from mock-treated and infected Obelisk plants, respectively. Dark and light green bars indicate the proteins isolated from mock-treated and infected Chinese Spring plants, respectively. (D) Western blot analysis for the abundance of the Rubisco large subunit was used as a marker for intra-cellular contamination of the apoplastic fluid samples. Histone H3 protein as a general eukaryote positive marker. (E) Colony morphologies grown on Petri dishes exemplify the microbial diversity recovered from the apoplastic fluids of each wheat genotype and treatment

**Figure S2. Contrasting susceptibility or resistance of wheat against *Zymoseptoria tritici* is due to the resistant gene Stb6 and the effector protein AvrStb6.** Representative asymptomatic leaves at 8 days post-inoculation (dpi) show the biotrophic phase of *Z. tritici* in both the susceptible Obelisk (without the resistant gene *Stb6*) and the resistant Chinese Spring (with the resistant gene *Stb6*) wheat plants. Representative leaves at 14 dpi show typical symptoms of *Z. tritici* infection in the susceptible Obelisk, including the chlorosis, necrosis, and the onset of asexual reproduction demonstrating by the pycnidial formation. On the other hand, Chinese Spring plants display a resistant phenotype due to Stb6-AvrStb6 interaction in this wheat genotype.

**Figure S3. Composition of the wheat-enriched endophytic community-based culture.** Wheat-enriched endophytic community-based culture collection is composed of 188 purified isolates which were classified taxonomically using partial 16S rRNA and ITS sequences. (A-C) Bar charts show the phyla-, classes- and families-level profiles obtained for the microbial diversity recovered from the apoplastic fluids of mock-treated and infected Obelisk and Chinese Spring wheat genotypes. (D-F) Venn diagrams showing the number of shared OTUs between groups and for the different taxonomical classifications (phylum, class, and family). Dashed black circles indicated the OTU commonly found among the treatments. (D) Proteobacteria and Actinobacteria were the commonly identified phyla. (E) Actinomycetia, Alphaproteobacteria and Gammaproteobacteria were the commonly identified classes. (F) Microbacteriaceae and Psedomonadaceae were the commonly identified families.

**Figure S4. Overview of antagonistic effect of the wheat-associated microbes against the Dutch IPO323 *Zymoseptoria tritici* isolate.** Species names are displayed in the bottom right corner (read from left to right). Microbes that did not grow are listed but were excluded from the data analysis. Three independent Petri dishes were performed per microbial endophyte.

**Figure S5. Overview of antagonistic effect of the wheat-associated microbes against the Danish Zt05 *Zymoseptoria tritici* isolate.** Species names are displayed in the bottom right corner (read from left to right). Microbes that did not grow are listed but were excluded from the data analysis. Three independent Petri dishes were performed per microbial endophyte.

**Figure S6. Overview of antagonistic effect of the wheat-associated microbes against the Iranian Zt10 *Zymoseptoria tritici* isolate.** Species names are displayed in the bottom right corner (read from left to right). Microbes that did not grow are listed but were excluded from the data analysis. Three independent Petri dishes were performed per microbial endophyte.

**Figure S7. Overview on the antagonistic activity of microbial organisms against three *Zymoseptoria tritici* strains and inhibition phenotypes.** (A) Venn diagram comparing the count of antagonistic microbes against the three different *Z. tritici* isolates in the confrontation assays. Each color refers to a different Z. tritici isolate, such as the Dutch IPO323, the Daish Zt05, and the Iranian Zt10. (B) Portions of gram-negative or -positive bacteria that inhibited *Z. tritici* isolates in the confrontation assays. Fisher’s exact tests were used to examine the association between the percentage of isolates inhibiting the *Z. tritici* isolate and the gram stain of the bacteria. (C) The phylogenetic tree was reconstructed based on partial 16S rRNA and ITS sequences. Nodes are supported based on 500 bootstraps performed in RAxML. Two fungal species *Rhizopus arrhizus* and *Mucor kunryamgriensis* from the Mucoroycota were used as outgroups in the tree. Antagonistic microbes and their interaction with the three tested *Z. tritici* isolates. The size of the circles is proportionate to the size of the inhibition zones. Values in the circle correspond to the average size in cm^2^ of the three technical replicates. Red circles represent the antagonistic microbes; white circles indicate the microbes that did not inhibit *Z. tritici*, and black circles demonstrated the microbes not tested in these antagonistic assays.

**Table S1.** Overview of the proteomics data of mock-treated and infected Obelisk and Chinese Spring wheat genotypes with *Zymoseptoria tritici* IPO323 strain.

**Table S2.** The differentially expressed proteins in the Obelisk and Chinese Spring wheat genotypes at 8 days-post incubation.

**Table S3.** The common 24 overexpressed during *Zymoseptoria tritici* IPO323 strain infection in Obelisk and Chinese Spring wheat genotypes.

**Table S4.** The most abundant Gene ontology (GO) enrichment terms for the upregulated proteins (mock vs infected) in Obelisk and Chinese Spring wheat genotypes.

**Table S5.** The most abundant Gene ontology (GO) enrichment terms for the downregulated proteins (mock vs infected) in Obelisk and Chinese Spring wheat genotypes.

**Table S6.** Proteomic analysis identified two up-regulated proteins and one down- regulated protein from Zymoseptoria tritici during plant-pathogen interaction.

**Table S7.** 188 microbes represent the wheat-enriched endophytic community-based culture collection used in this study.

**Table S8.** Frequency of each phenotype within five time points and total phenotype frequency from the microtiter plate experiment.

**Table S9.** Overview on select top 10 bacterial strains with the highest and the lowest OD ratio per plant compound within each time point.

