## Supplementary figures and images for "The apoplastic space of two wheat genotypes provide highly different environment for pathogen colonization: Insights from proteome and microbiome profiling"

### Supplemental Figure 1

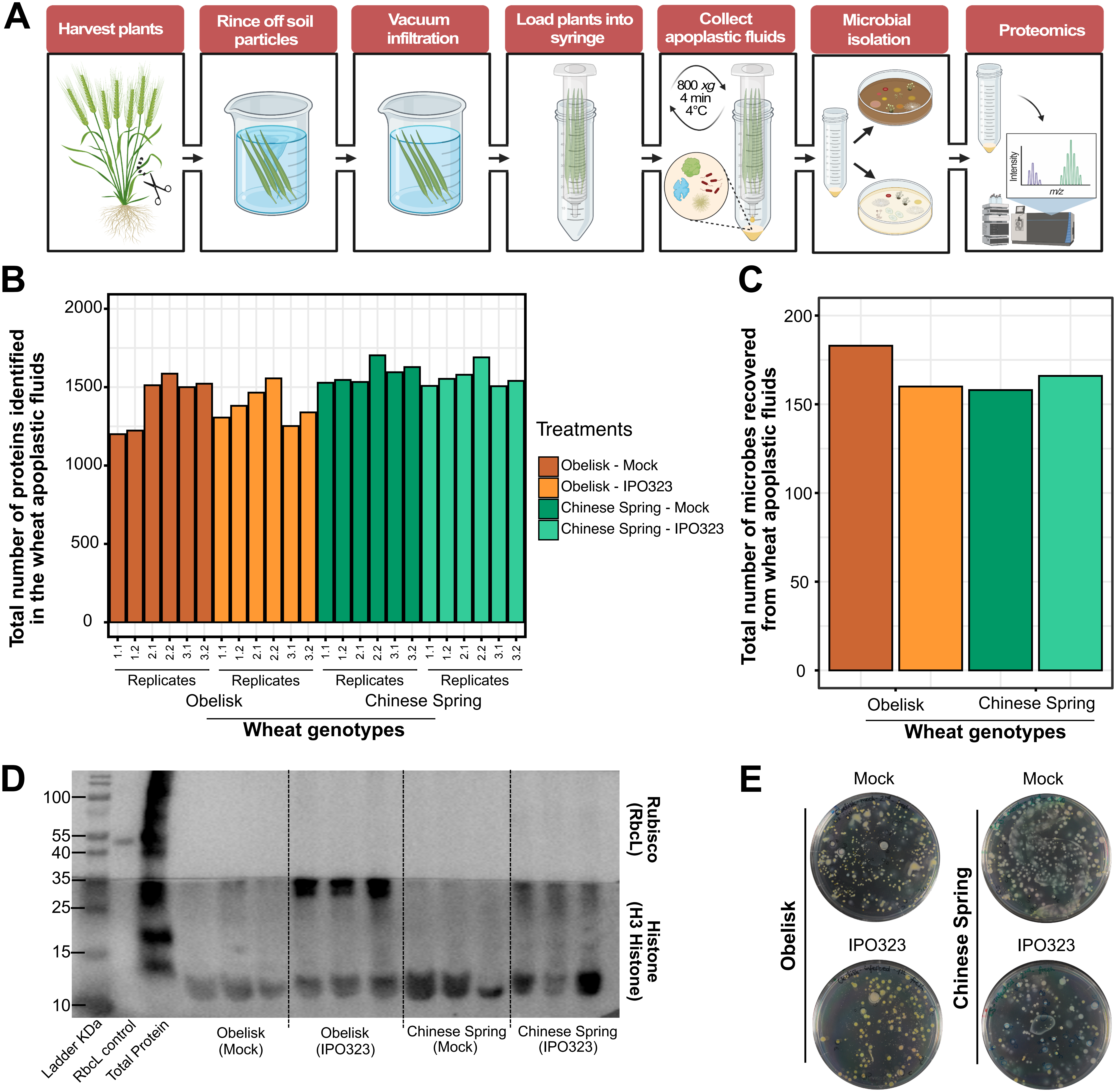

### Supplemental Figure 2

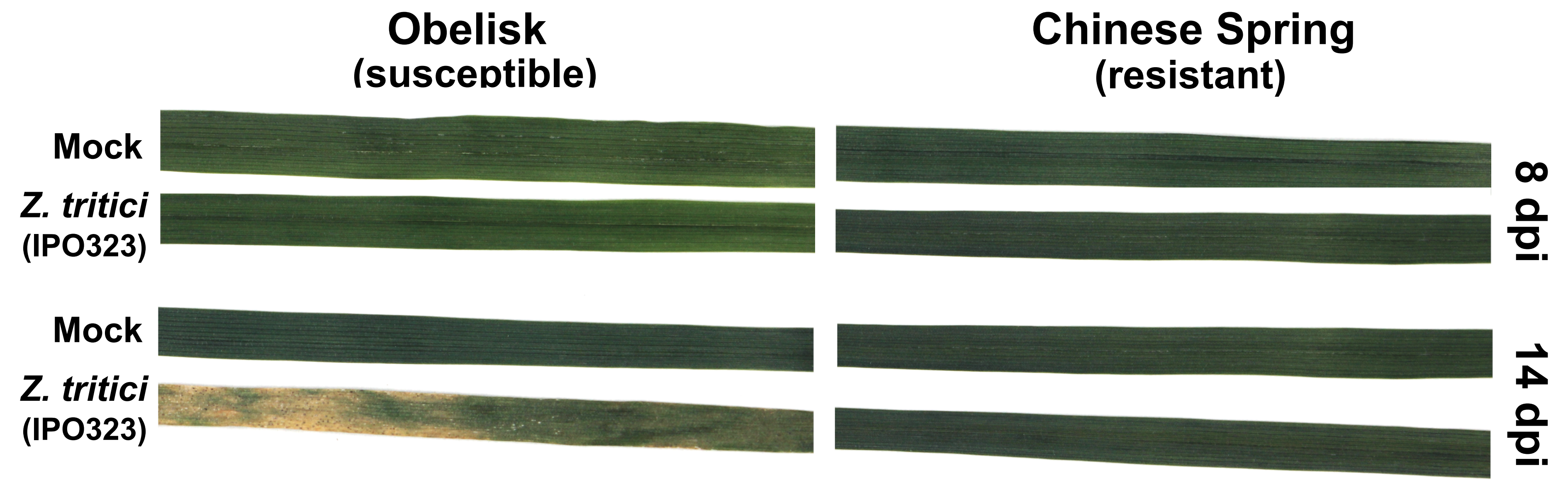

### Supplemental Figure 3

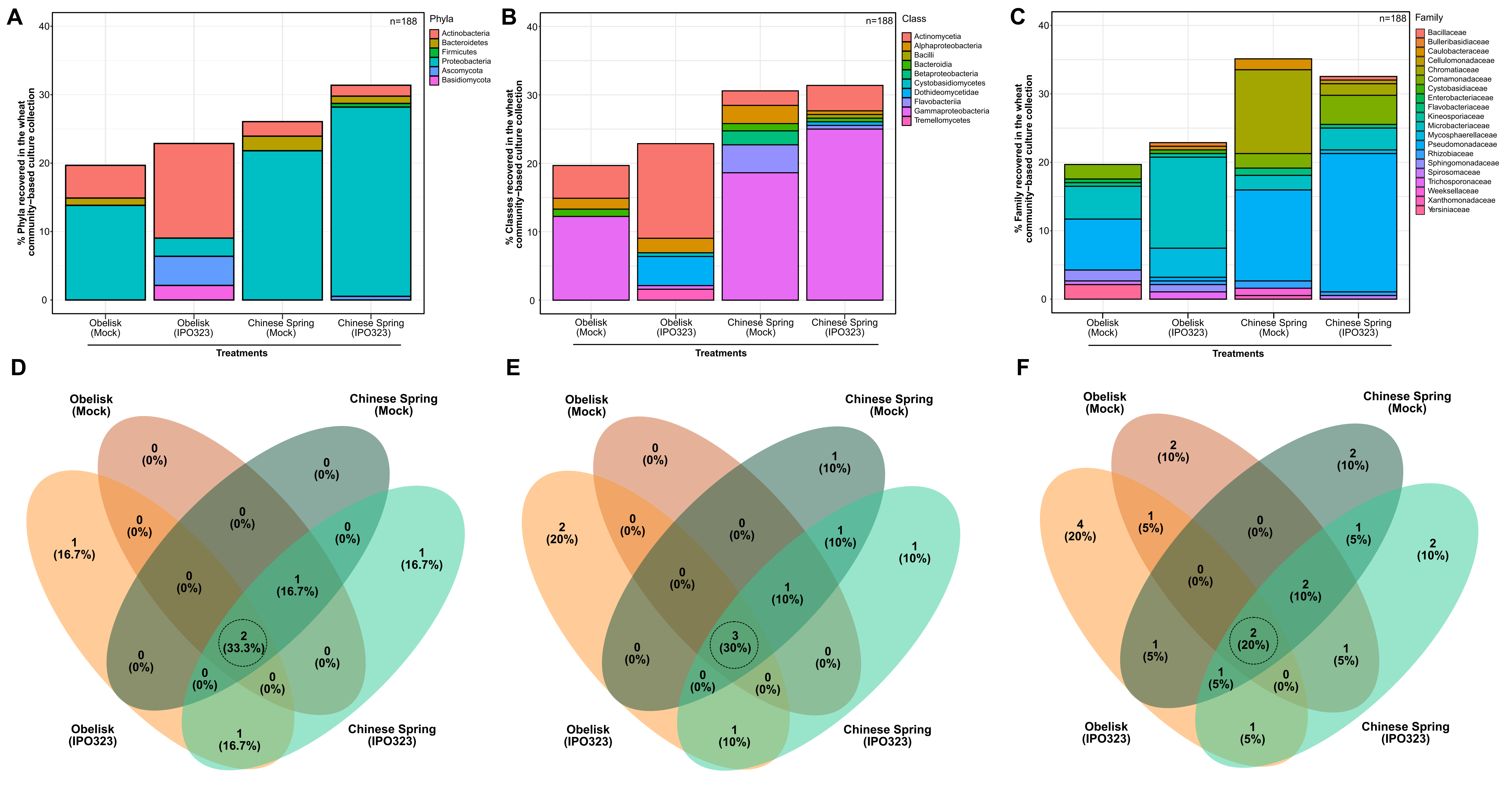

### Supplemental Figure 4

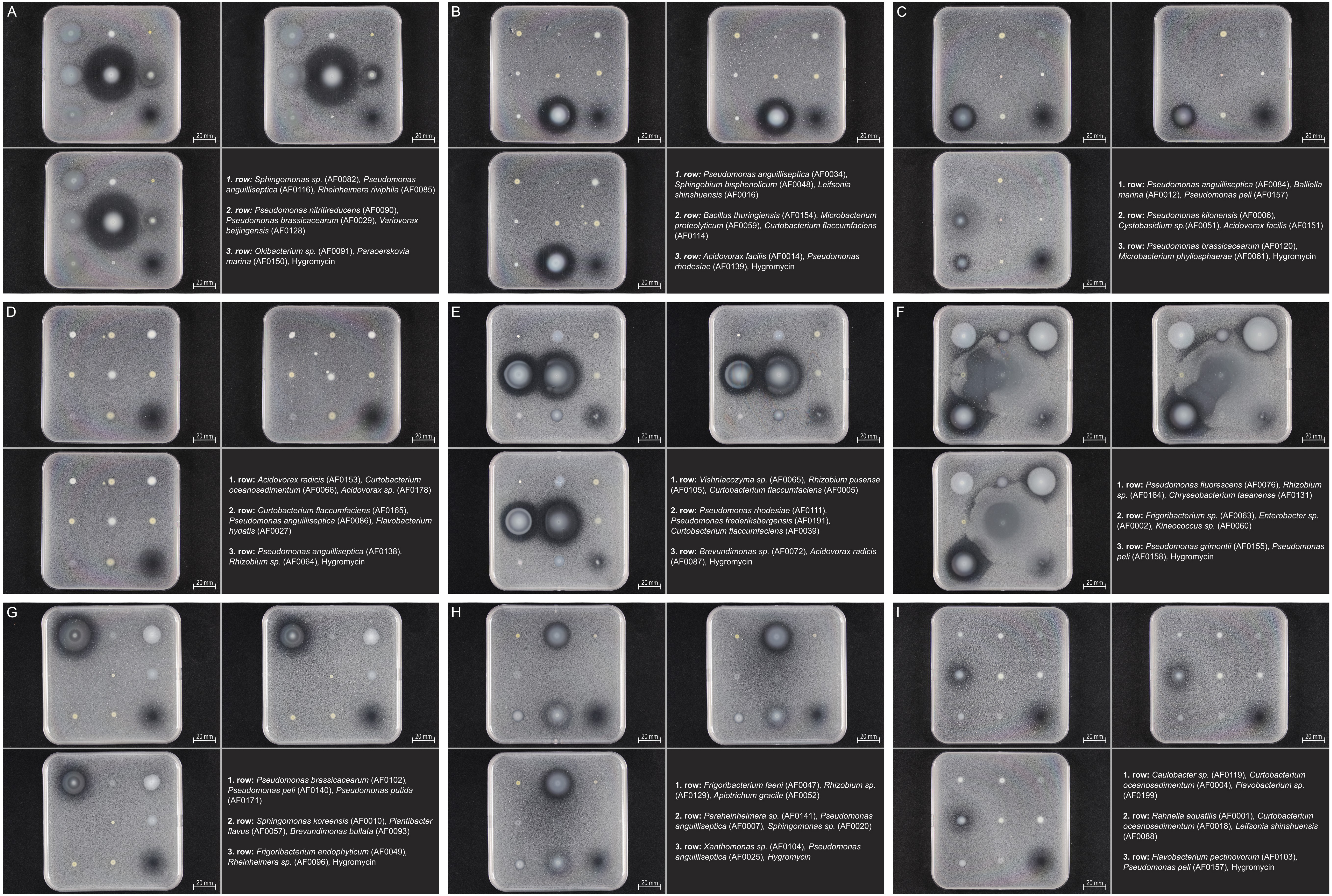

### Supplemental Figure 5

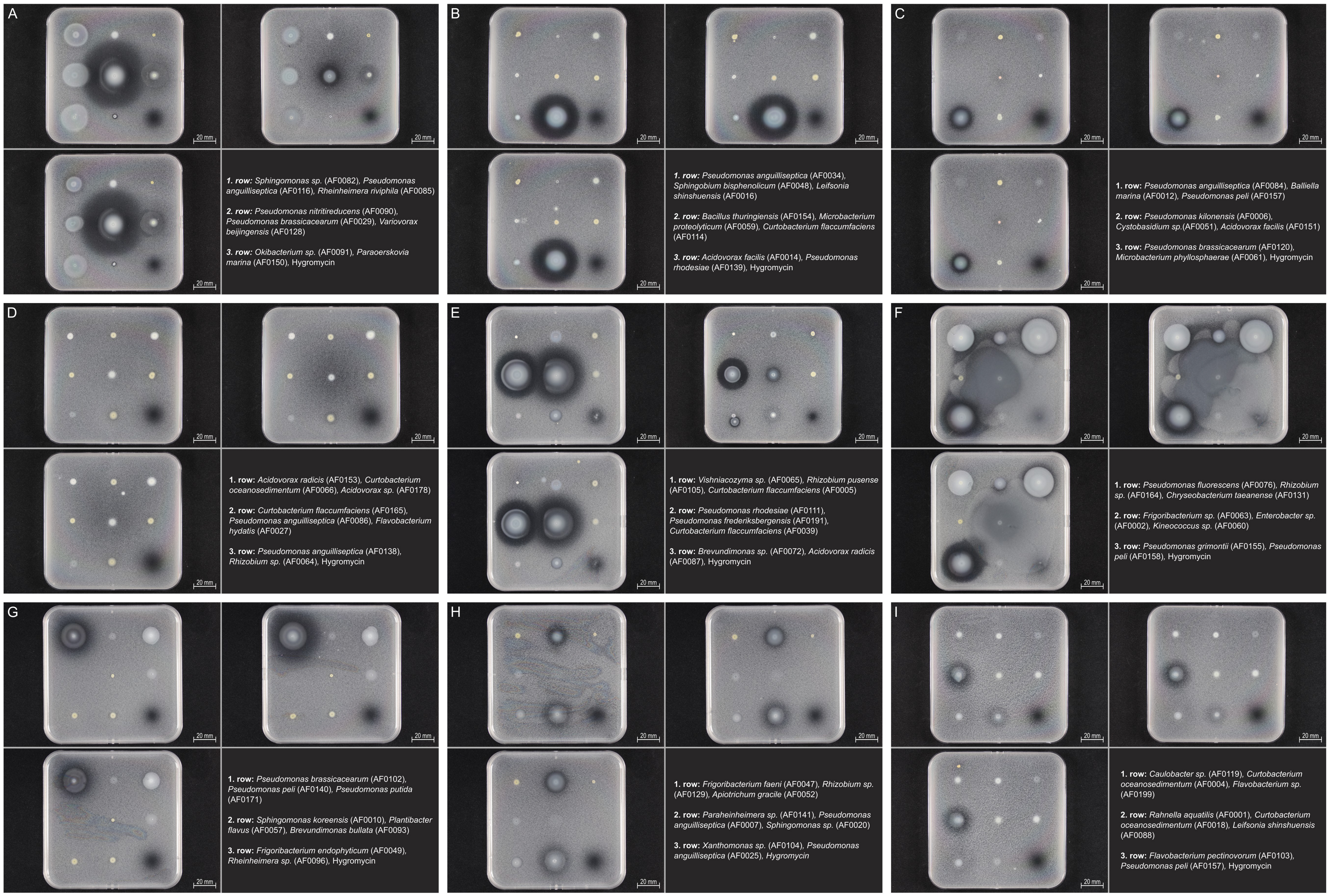

### Supplemental Figure 6

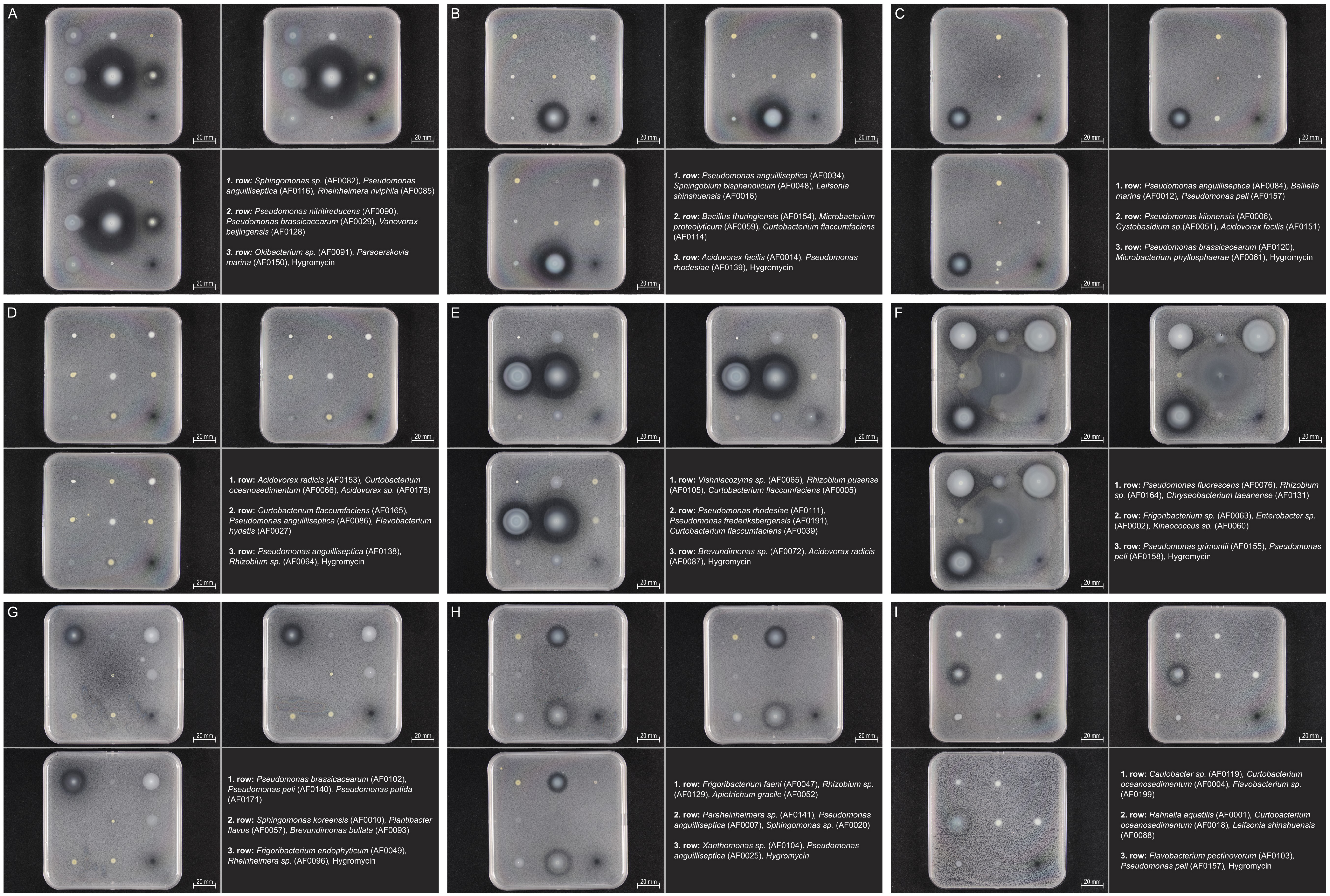

### Supplemental Figure 7

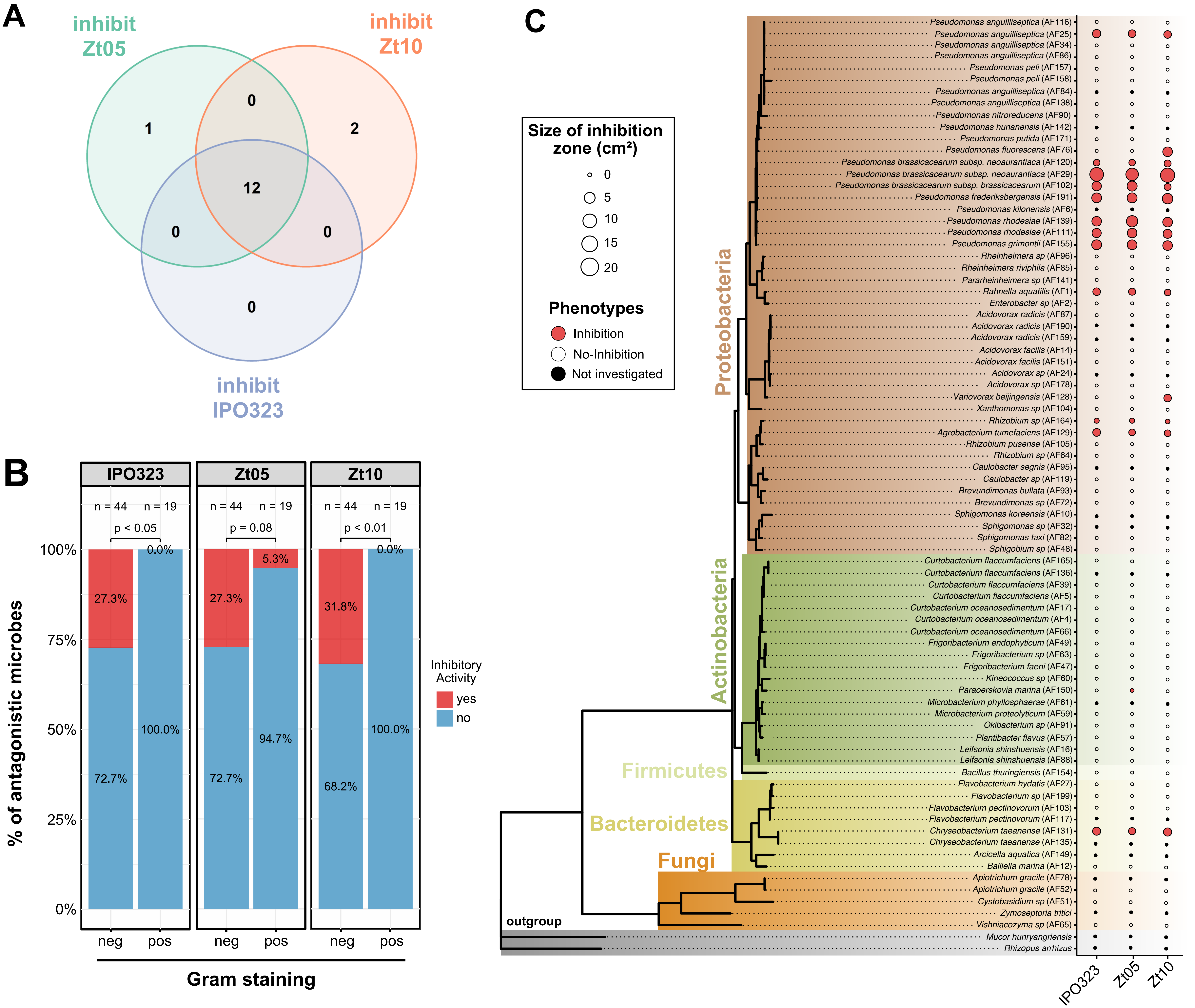
